# Maternal senescence broadly reprograms gene expression in offspring

**DOI:** 10.64898/2026.04.28.720237

**Authors:** Soleille Morelli Miller, Zachariah Wylde, Russell Bonduriansky

## Abstract

Offspring of older parents have reduced fitness in many species, but the mechanisms mediating this cross-generational dimension of ageing remain poorly understood. Senescence is associated with genome-wide epigenetic changes that alter transcription, raising the possibility that older parents transmit dysregulated gene expression patterns. Here we show that maternal senescence induces deleterious, transcriptome-wide reprogramming of gene expression in offspring. Gene ontology and pathway enrichment analyses reveal that broad changes in gene expression that characterise maternal senescence in the clonally reproducing arthropod *Folsomia candida* are also observed at a young age in the offspring of older mothers. These concordant fold-changes are apparent at the whole-transcriptome level, and encompass conserved sequalae of senescence, including reduced carbon and energy metabolism. However, senescent mothers and their offspring also exhibit some contrasting gene expression patterns, representing distinct transcriptomic signatures of senescence that are not reflected in its cross-generational effects. The broad reprogramming of gene expression in offspring of older mothers is associated with substantially reduced fitness. Our findings show that older females can transmit a senescence-like gene expression syndrome to their offspring, inducing deleterious phenotypes from a young age.

## Introduction

As organisms age, cellular processes deteriorate throughout the body (Lu et al. 2026), leading to a global decline in physiological function that typically drives increased mortality risk and reduced reproductive performance (Jones et al. 2014). Beyond effects of senescence on fertility (Serre and Robaire 1998; Andersen 2000) and mating success (Bonduriansky and Brassil 2002; Ruhmann et al. 2018), parental ageing can also shape offspring phenotypes (Lansing 1947, 1954; Monaghan et al. 2020). Such “Lansing” or “parental age” effects have been documented across diverse taxa, often resulting in developmental abnormalities (Pellestor et al. 2003; Keefe et al. 2005; Wu et al. 2017) as well as reduced pre-adult survival, adult lifespan, and reproductive success (Amarya et al. 2018; Ivimey-Cook and Moorad 2020) in offspring of older parents (although neutral or even beneficial outcomes have also been reported (Ivimey-Cook and Moorad 2020; Monaghan et al. 2020)). This cross-generational dimension of senescence could represent an important source of variation in fitness, and potentially influence the evolution of reproductive strategies and life histories (Priest et al. 2002; Wylde et al. 2019). From a public health perspective, the increasing trend toward delayed motherhood in many human populations (Burkimsher 2015; Bongaarts et al. 2017; Oecd 2024) also highlights the importance of understanding the biological mechanisms mediating parental age effects.

Several conserved cytological hallmarks of aging have been identified across diverse animal taxa (López-Otín et al. 2013; Frenk and Houseley 2018; López-Otín et al. 2023). These cytological hallmarks include widespread downregulation of genes encoding mitochondrial proteins (e.g. components of the electron transport chain (De Magalhães et al. 2009; Glass et al. 2013; Kumar et al. 2013; Van Den Akker et al. 2014; Peters et al. 2015; Ma et al. 2016; Cannon et al. 2017; Webb and Sideris 2020)), protein synthesis and recycling (e.g. ribosomal proteins and ribosome biogenesis factors (Berchtold et al. 2008; Philipp et al. 2013; Kamei et al. 2014; Janssens et al. 2015; Bryois et al. 2017; Choi et al. 2024)), and growth factors (Zahn et al. 2005; Schumacher et al. 2008; Ma et al. 2016). Aging is also associated with the dysregulation of immune system functions, typically involving an overexpression of pro-inflammatory mediators and upregulation of immune response genes (e.g. IL-1β, IL-6, COX-2, TNF-α, Casp1)(Kim et al. 2002; Terao et al. 2002; Chung et al. 2006; De Magalhães et al. 2009; Mirza et al. 2011; Lee et al. 2012; Carlson et al. 2015; Bartling et al. 2019), and a decline in neural function (Burke and Barnes 2006). The typically deleterious effects of advanced parental age at reproduction on offspring suggest that some age-related physiological changes are transmitted across generations. Yet, the mechanisms mediating the effects of parental age on offspring remain poorly understood.

Senescence is associated with extensive changes in gene expression across the genome, potentially reflecting age-related epigenetic dysregulation (Bollati et al. 2009; Horvath et al. 2012; Horvath 2013; Horvath and Raj 2018; Teschendorff and Horvath 2025; Lu et al. 2026). Such age-related changes (represented by “epigenetic clocks”) encompass the parental germ-line epigenome (Kawai et al. 2018; Perez and Lehner 2019), raising the possibility that older parents transmit senescent patterns of gene expression to their offspring and thereby inducing deleterious phenotypes in offspring from a young age (Bonduriansky and Day 2018; Ashapkin et al. 2023). Indeed, human cohort studies show that maternal age is associated with patterns of DNA methylation at some loci in children (Jiang et al. 2024; Yeung et al. 2024), and affects gene expression in human oocytes (Steuerwald et al. 2007; Ntostis et al. 2022). Moreover, some genes show similar patterns of DNA methylation and expression in aged parents and their children (Hua et al. 2022). However, the potential for broad transmission of senescent gene expression patterns to offspring has not been investigated experimentally. In rodents, offspring of aged parents exhibit altered DNA methylation and gene expression in the embryonic brain (Duan et al. 2015; Yoshizaki et al. 2021), but the transcriptomic breadth and pattern of reprogramming remain unclear. Importantly, it is not known to what extent parental age effects are manifested in concordant patterns of gene expression across generations *versus* distinct gene expression profiles in senescent parents and their offspring. We define concordance of gene expression between generations as a positive correlation between fold-changes in old *versus* young parents (ΔF0) and fold-changes in offspring (assayed at a young age) of old *versus* young parents (ΔF1). Experimental evidence of broad concordance would support locus-specific transmission of senescent gene expression across generations. Here, we utilized the clonally reproducing arthropod *Folsomia candida* (Fountain and Hopkin 2005) to investigate effects of maternal age on offspring gene expression and fitness within multiple isogenic lineages.

## Methods

### Lab colonies

*Folsomia candida* is a collembolan arthropod (“springtail”) that reproduces clonally, allowing for the establishment of an isogenic line (isoline) from a single female (Fountain and Hopkin 2005). Clonal reproduction makes it possible to investigate effects of maternal age on offspring while avoiding the potentially confounding effects of selection on genetic variation and the complications introduced by recombination and sexual reproduction. Ten isolines were established in 2023 from single females derived from our laboratory-reared stock population (fed on yeast provided on a substrate of cow dung and cocopeat). This stock originated from individuals from a wild population collected by Dr. Penelope Greenslade in the Jenolan Caves, NSW, Australia, and was maintained in our laboratory from 2020.

Before creating the experimental populations for this study, a standardization step was implemented as follows: Eggs were collected from each isoline. Once these eggs hatched and nymphs reached 7–10 days old (large enough to not be damaged during aspirator transfer), they were standardized to a density of ∼50 individuals per container. When these populations reached 20–25 days old, offspring from this generation were used to establish the parental generations for the subsequent experiments described below.

### Animal husbandry

All springtails used in these experiments were housed in 250 mL containers on a substrate (ca. 1 cm depth) of plaster of paris (3201, Uni-Pro Pty Ltd, QLD, Australia) and activated charcoal (CL026, ChemSupply Australia Pty Ltd, SA, Australia) according to OECD guidelines(Oecd 2016) for rearing of *F. candida*. Springtail housings were kept at 25 (± .5) ºC and 100% RH (substrate saturated with RO water) in complete darkness. All adults were fed *ad libitum* from one batch of dried baker’s yeast (*Sacchromyces servisiae*) (Nixon bulk foods, Australia) that was stored at -20 ºC until required. Housings were renewed weekly to avoid the proliferation of other fungi and accumulation of waste products, and to keep population densities constant. Adult springtails were transferred using an aspirator, and eggs were carefully placed onto black filter paper (Westlab, 409009) using a moistened paintbrush. Population densities were standardised 7-10 days after hatching when individuals were large enough to be transferred without causing damage.

### Experimental design

We aimed to test whether maternal age can imprint offspring transcription profiles, such that old mothers produce offspring with a senescence-like transcription profile from a young age (Fig. 1). We also examined age-related differences in fitness-related phenotypes between offspring produced by young versus older mothers. This project was pre-registered with the Open Science Framework (10.17605/OSF.IO/H87BU). Data and code are available from the Zenodo repository (Miller et al. 2026).

**Figure 1.**
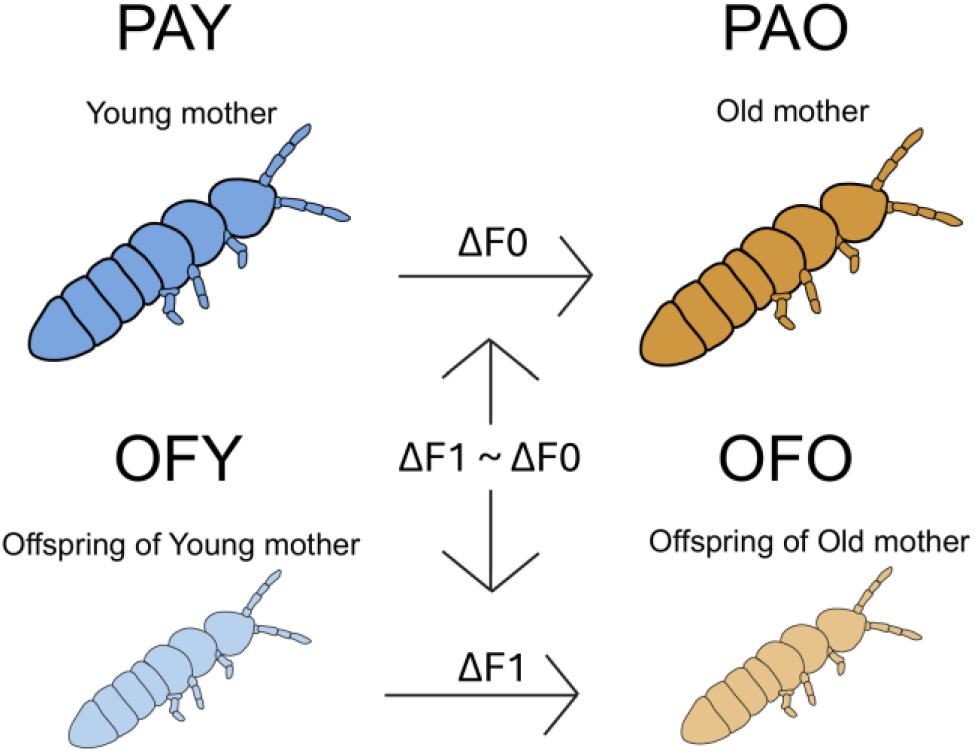
Experimental design: Young mothers (PAY) and older mothers (PAO) produced OFY and OFO offspring, respectively. We then quantified differential gene expression between PAO and PAY adults (ΔF0); differential gene expression between OFO and OFY offspring, all aged 20-25 days (ΔF1); and the correlation of fold-change across generations (ΔF1 ∼ ΔF0).

For gene expression analysis, we established six replicate F0 populations from each of 10 isolines. Each replicate population consisted of a clone produced by a single founder female nymph, and all replicate populations consisted of 30 individuals per population. From three replicate F0 populations per isoline, eggs were collected at age 20-25 days (young maternal age at breeding: PAY) to generate F1 offspring (OFY). From another three replicate F0 populations per isoline, eggs were collected at age 70-75 days (older maternal age at breeding: PAO) to generate F1 offspring (OFO). Immediately after egg collection, all F0 replicate populations were frozen in liquid nitrogen and then stored at -80 ºC. Offspring (F1) from each of the replicate populations were kept at the same density and environmental conditions as their mothers and then harvested at age 20-25 days (n = 40 offspring per replicate population) for RNAseq. Additional individuals from each replicate population were harvested for morphometric analysis. All F1 samples were immediately frozen in liquid nitrogen and then stored at -80 ºC until RNA extraction. One isoline (isoline 3) was excluded from analysis because of poor RNA yield.

### Reproduction and survival

The same 10 isolines were used to establish another twenty replicate populations, each replicate population consisting of five same-age individuals. From 10 replicate F0 populations per isoline, eggs were collected at age 20-25 days (PAY treatment) to obtain F1 offspring (OFY). From the other 10 replicate populations per isoline, eggs were collected at age 40-45 days (PAO treatment) to obtain F1 offspring (OFO). OFO eggs were thus harvested at younger maternal age for reproduction and mortality analysis than for gene expression analysis, providing a conservative assessment of the phenotypic effects of maternal senescence on offspring fitness.

For each replicate population, development time was quantified as the number of days from eggs laid to first F1hatching, and age at first reproduction was quantified as the number of days from first F1 hatchling to first F2 eggs laid. The number of live F1 adults was recorded weekly to estimate mortality rate. To minimize build-up of waste products, populations were provided with a substrate composed of plaster mixed with activated charcoal and each replicate population was transferred to a fresh container weekly. The 20 F1 populations (OFY and OFO) were maintained, and their fecundity quantified weekly (see below), until at least 3 of the 5 individuals in the population had died or the population reached 167 days of age.

To estimate age-specific fecundity, each F1 replicate population container was photographed at least once per week from above using a Sony RX10 camera and new eggs or hatchlings were counted, resulting in a total of 2,210 weekly fecundity estimates comprising 274,493 F2 offspring. For practical reasons, fecundity was estimated from a combination of egg and hatchling counts, depending on the stage of F2 development at each observation. Combining egg and hatchling counts is unlikely to introduce bias because hatching success was typically > 80% and egg/hatchling counts were randomly distributed among replicates and ages. All egg counts were done manually, but some hatchling counts were done using automated image analysis. For automated counts, images were automatically cropped to the plaster surface area using a circular Hough transform and subdivided into overlapping tiles (640 × 640 pixels, 25% overlap) prior to analysis. A YOLOv8 object detection model implemented using the Ultralytics framework (Jocker et al. 2023) was trained to detect individual hatchlings using 470 manually annotated image tiles containing bounding boxes for each hatchling, together with 30 background tiles containing no hatchlings. Of these, 400 tiles were used for model training and 100 were withheld for internal validation during training. Models were trained for 40 epochs. During inference, detections were generated at an image resolution of 640 pixels and filtered using a confidence threshold of 0.35 and non-maximum suppression with an intersection-over-union threshold of 0.5. Predictions from overlapping tiles were recombined to produce hatchling counts for each container image. Model performance was evaluated using the independent validation set of 100 container images not used during model training. Automated counts showed strong agreement (*r* = 0.99) with manual counts (see Figure S4). For images containing hatchlings, oviposition dates were estimated based on a temperature-specific mean egg development time of 7 days at 25 °C. Code, trained model weights, and image-processing scripts are available on the Github repository (https://github.com/wyldescience/FolSum).

### RNA isolation, sequencing, and quality control

From each replicate population, forty springtails were included in each cryotube as a pooled sample. Total RNA was extracted using QIAzol® Lysis Reagent (QIAGEN®, Hilden, Germany) following the manufacturer’s protocol. DNase treatment (DNase I Solution, Thermo Scientific™) was then performed in a thermocycler per the manufacturer’s instructions. The samples were washed with ethanol to remove the remaining DNAse. Samples were tested for quantity and quality with a NanoDrop® ND-1000 (Thermo Scientific™) and a Qubit™ 4 Fluorometer (Invitrogen, Thermo Fisher Scientific, Singapore) with the Broad-Range RNA Assay Kit. Samples were stored at -80 ºC until sequencing.

A total of 64 purified total RNA samples, comprising all combinations of isoline × treatment × generation, with some combinations represented by two replicate populations (see Table S2), were submitted to the Ramaciotti Centre for Genomics at UNSW Sydney, Australia, for library preparation and sequencing. Stranded paired-end RNA-seq libraries were generated and sequenced using a NovaSeq X Plus system with an S4 flowcell, producing 2 × 100 bp paired-end reads. After library preparation and sequencing, quality control of the raw sequencing data was performed using FastQC version 0.11.9 (Andrew 2010). To assess the impact of trimming, we tested the effect of removing 10 bp from the 5’ end and 2 bp from the 3’ end due to slight bias in per base sequence content. However, untrimmed reads exhibited higher mapping rates to the reference genome, and no (or modest) trimming results in the most biologically accurate gene expression estimates (Williams et al. 2016; Jocker et al. 2023), so we proceeded without trimming. Untrimmed paired-end reads were aligned to the *Folsomia candida* reference genome and transcriptome (GenBank accession: GCF_002217175.1) using STAR aligner version 2.7.11b (Dobin et al. 2013) with default parameters. A total of 2,829,509,816 reads were obtained from 64 samples with an average of 44 million reads per sample. Unique mapping rates to the *Folsomia candida* reference genome ranged from 70.45% to 85.92% (mean = 79.60%; see Table S2). The resulting BAM files were processed with *featureCounts* from subread v.2.0.2 (Liao et al., 2014) to generate unnormalized gene-level read count matrices. Replicate populations were collapsed to generate 36 groups (9 isolines × 2 generations × 2 treatments) for differential expression analysis.

### Variability of gene expression

We first investigated whether age influenced the variability of gene expression profiles. According to the mutation accumulation theory of senescence (Medawar 1952; Hughes et al. 2002), selection strength becomes weaker as a cohort ages, allowing mutations with late-life expression to accumulate and resulting in greater transcriptional variability. To test this, we assessed gene expression variability across treatment groups in both the parental (PAO vs. PAY) and offspring (OFO vs. OFY) generations. First, we visualized variation in gene expression using principal component analysis (PCA) implemented via the *prcomp()* function from the *stats* package (version 3.6.2) in R. We then tested for differences in expression variability using a multivariate homogeneity of group dispersions test using *betadisper()* from the *vegan* package (version 2.6-1 (Dixon 2003)), applied to the normalized (rlog-transformed) count matrix.

### Differential gene expression and functional enrichment analyses

To identify differentially expressed genes (DEGs) between older and young F0 adults (PAO vs. PAY, denoted ΔF0; Figure 1) and between their F1 offspring at age 20-25 days (OFO vs. OFY, denoted ΔF1; Figure 1), we performed two separate genome-wide differential expression analyses using *DESeq2* (version 1.48.1; Love et al., 2014). *DESeq2* accounts for multiple testing by applying the Benjamini–Hochberg correction to control the false discovery rate (FDR), generating adjusted *p*-values. Genes with an adjusted *p*-value < 0.05 and an absolute log_2_ fold change > 1 were considered DEGs.

DEGs were functionally annotated with Gene Ontology (GO) terms using the orthology-based annotation tool eggNOG-mapper (emapper version 2.1.12; http://eggnog-mapper.embl.de) with default settings. Although a greater proportion of DEGs could be annotated using the broader Eukaryotic database (see Supplementary Information), we prioritised Arthropoda-based one-to-one ortholog annotations as they provide a more phylogenetically relevant functional context for springtail genes. This taxon-restricted approach reduces potential annotation noise from distant homologs and improves the reliability of downstream GO-based interpretation. KEGG pathway annotations were retrieved for *Folsomia candida* from the *AnnotationHub* resource (accession=AH115244). Functional enrichment analyses were conducted for both comparisons (PAO vs PAY and OFO vs OFY) in R (version 4.5.0) using the enricher function for GO term enrichment, enrichKEGG for KEGG pathway enrichment, and gseKEGG for gene set enrichment analyses (GSEA) from the package *clusterProfiler* (version 4.16.0).

### Gene expression differences across generations

To test whether age-related gene expression changes in mothers (F0) were reflected in their offspring (F1), we first identified genes that were upregulated or downregulated in both generations (i.e., showing the same direction of fold-change in ΔF0 and ΔF1), and genes showing opposing regulation in F0 vs. F1 generations (i.e., showing opposite directions of fold-change in ΔF0 versus ΔF1). We used *ggVennDiagram* to visualize the overlap of differentially expressed genes (DEGs) between generations.

We then used the *rstatix* package (v0.7.2) to calculate Pearson’s correlation coefficient between the log_2_ fold changes in older versus young mothers (ΔF0) and the log_2_ fold changes in offspring of older versus young mothers (ΔF1). We quantified the fold-change correlation (ΔF1 ∼ ΔF0) separately across all genes (n=22,555), as well as genes upregulated in both generations (*n* = 19), genes downregulated in both generations (*n* = 112), genes upregulated in older mothers but downregulated in their offspring (*n* = 7), and genes downregulated in older mothers but upregulated in their offspring (*n* = 30).

To determine the probability that these ΔF1 ∼ ΔF0 correlations occurred by chance or as an artefact of our design, we performed a randomization test to generate null distributions for comparison. Specifically, we simulated gene expression count matrices for each of the four experimental groups (PAY, PAO, OFY, OFO) using a negative binomial distribution, which is well-suited for bulk RNA-seq data due to its ability to model overdispersion, which is typically present in RNA-seq count data (Robinson et al. 2010). For each gene, simulated counts were constrained by the observed overall mean and variance of that gene in the real dataset. Using these simulated matrices, we performed differential gene expression analyses for ΔF1 and ΔF2, following the same pipeline as used for the observed data. From each simulation, we recorded the number of DEGs unique to each generation, and the number shared across both generations. This process was repeated 10,000 times to generate null distributions. We then compared these simulated values to the observed values. We also used the simulated data to quantify ΔF2 ∼ ΔF1 correlations as described above to generate a null distribution for the correlation coefficients (see Simulation Analysis in Supplementary Information for further details).

### Reproduction analysis

To investigate effects of maternal (F0) age at reproduction on offspring (F1) fitness-related phenotypes, linear mixed models were fitted in R version 4.4.2 using the package *glmmTMB* version 1.1.14 (McGillycuddy et al. 2025). Where required, heterogeneity of variances was accounted for using the function *dispformula* and zero-inflation was accounted for using the function *ziformula*. Model assumptions were tested using the packages *DHARMa* version 0.5.0 (Hartig 2025) and *performance* version 0.16.0 (Lüdecke et al. 2021). F1 development time (days from oviposition to first hatchling for each replicate population) was analysed using a mixed model with maternal (F0) age at reproduction (OFO vs. OFY) as a fixed, categorical factor, isoline identity as a random factor, and Gaussian error structure. F1 age at first reproduction was analysed using a mixed model with maternal (F0) age at reproduction (OFO vs. OFY) as a fixed, categorical factor, isoline identity as a random factor, and negative binomial exponent error structure (with one very high value excluded to improve the distribution of residuals). Total fecundity of F1 replicate populations (up to age 167 days) was analysed using a mixed model with maternal (F0) age at reproduction (OFO vs. OFY) as a fixed, categorical factor, isoline identity as a random factor, and negative binomial exponent error structure. The total fecundity model was then re-fitted with reproductive lifespan (days between first reproduction and last reproduction for each F1 replicate) and mean number of individuals alive throughout the reproductive lifespan of each F1 replicate as additional fixed continuous predictors in order to estimate effects on individual daily reproductive rate. F1 age-specific fecundity was summed across 24-day age intervals and the number of eggs laid per age interval was then modelled with maternal (F0) age at reproduction (OFO vs. OFY) as a fixed categorical predictor and the mean number of individuals alive during the time interval and replicate population age at the end of the time interval as fixed continuous predictors, isoline identity and replicate population identity as random factors, and negative binomial error structure. To test overall effects of replicate population age and its interaction with maternal age at reproduction, we carried out a likelihood ratio test comparing the full model and a simplified model lacking the offspring age x maternal age interaction using the *anova* function from the *car* package version 3.1-5 (Fox and Weisberg 2019). For detailed statistical results, see the Supplementary Information (Tables S5 – S10).

### Survival analysis

Survival analysis was carried out using the *coxme* package version 2.2-22 (Therneau 2024), with maternal (F0) age at breeding (OFO vs. OFY) as the fixed effect and isoline identity as the random effect. Replicate F1 populations were treated as the unit of replication, with population lifespan defined as the number of days from hatching until most (i.e., at least 3) individuals had died. Populations surviving longer than 167 days were treated as right-censored data (Table S11).

### Body size analysis

Body size of F0 mothers (PAO and PAY) and F1 offspring (OFO and OFY) was quantified by mounting 2 to 5 individuals per replicate population in glycerol on microscope slides with cover-slips held ∼ 0.7 mm above the slide surface to avoid crushing the samples. Each individual was imaged using a Leica MC170 HD camera mounted on a Leica MZ 16A stereoscope, and total body length was quantified using ImageJ software(Schneider et al. 2012) from the images (Figure S3). Body length was analysed using a mixed model with treatment (PAO, PAY, OFO, OFY) as a fixed, categorical predictor, isoline identity and replicate population identity as random effects, and gaussian error structure (Table S12). Tukey post-hoc tests were used to compare groups (Table S13).

## Results

### Ageing alters gene expression in females

Within the maternal (F0) generation, A total of 812 genes showed significant differential expression in older females (age 70-75 days; PAO) when compared to young adult females (age 20-25 days; PAY), representing the transcriptional changes associated with senescence (ΔF0). Of the identified differentially expressed genes (DEGs), 25.6% (208) were upregulated and 74.4% (604) were downregulated in PAO compared to PAY (see Figure 2c). We also observed a trend toward reduced variability in gene expression profiles in PAO relative to PAY (multivariate homogeneity of group dispersions test, *p* = 0.06; Figure 2a).

**Figure 2.**
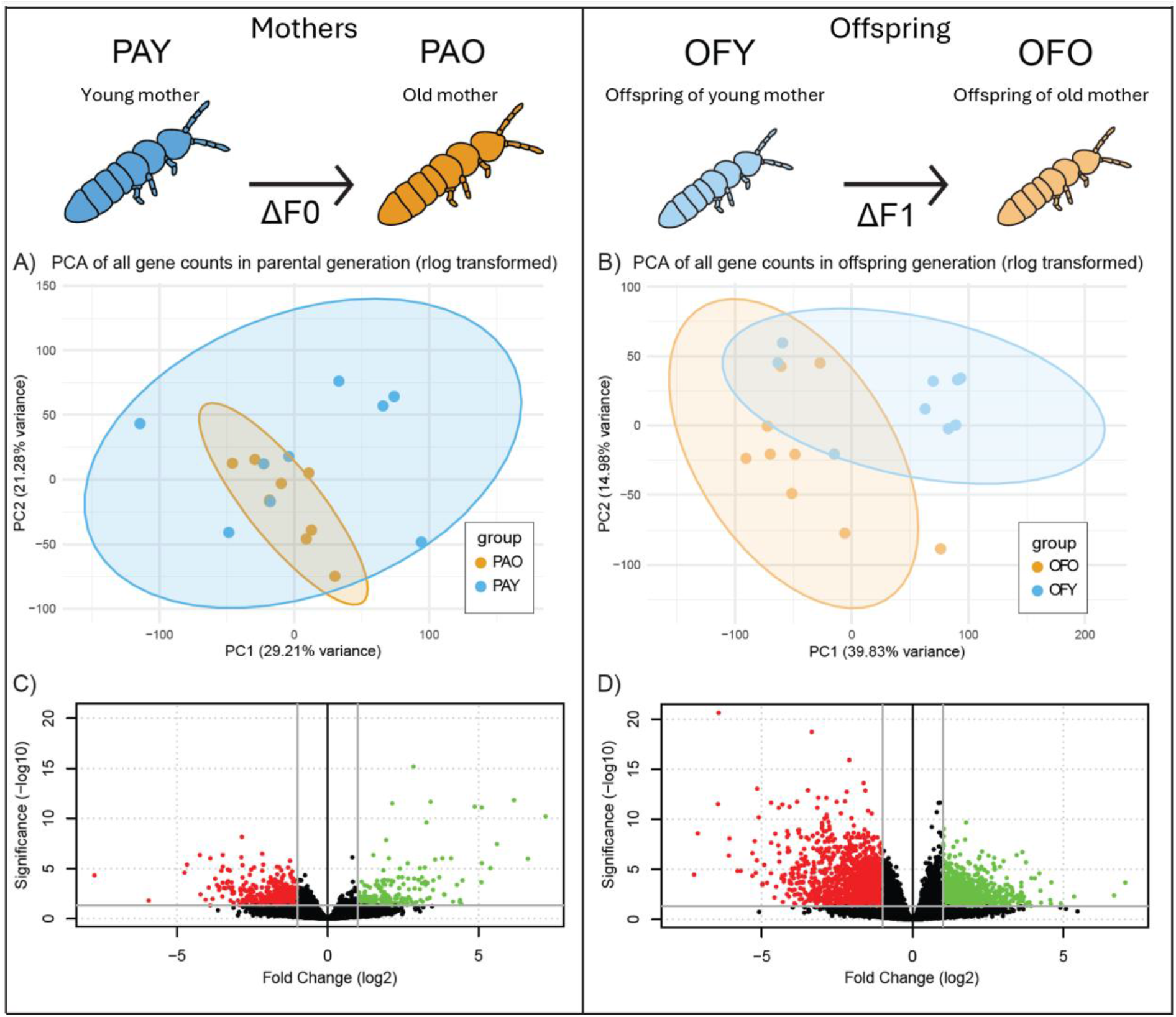
Gene expression profiles and differentially expressed genes in the parental (F0) and offspring (F1) generation. Principal component analysis (PCA) of rlog-transformed gene expression data in the parental generation (A) and offspring generation (B). Volcano plot showing differentially expressed genes (DEGs) between young females (PAY) and older females (PAO) in the parental (F0) generation, ΔF0 (C), and differentially expressed genes between offspring of older females (OFO) and offspring of young females (OFY) in the offspring (F1) generation, ΔF1 (D). Coloured points represent significantly upregulated (green), downregulated (red), and non-significant (black) genes in ΔF0 and ΔF1.

### Transcriptomic signatures of ageing

The transcriptional patterns we observed in older mothers, ΔF0, are consistent with recognised molecular hallmarks of ageing (López-Otín et al. 2013; Frenk and Houseley 2018). Of the 812 differentially expressed genes in ΔF0, 10.7% (87 genes, 7 upregulated, 80 downregulated) were able to be annotated with one-to-one orthologs in Arthropoda and these genes had GO terms assigned.

Ten GO terms were significantly enriched among downregulated genes in PAO (adjusted *p* < 0.05). The most highly enriched pathways were related to neural activity and synaptic signalling, including chemical and trans-synaptic transmission, larval behaviour, and nervous system processes (Figure 3a). Reduced neural plasticity and synaptic efficiency are recognised hallmarks of neurobiological ageing and have been linked to cognitive decline and slower behavioural responses in model organisms(Burke and Barnes 2006). In addition to these neural functions, broader biological processes such as cell–cell signalling and system processes were also downregulated, suggesting a general decline in neural activity and physiological signalling capacity in older adults. We also found 14 GO terms that were significantly upregulated in PAO compared to PAY (adjusted *p* < 0.05; Figure 3b). These included small molecule and lipid metabolic processes, as well as multiple pathways involved in hormone and steroid metabolism such as steroid, ketone, and ecdysteroid metabolic and biosynthetic processes. These upregulated GO terms suggest an age-associated increase in some metabolic and biosynthetic processes, particularly within hormone- and lipid-related pathways.

**Figure 2.**
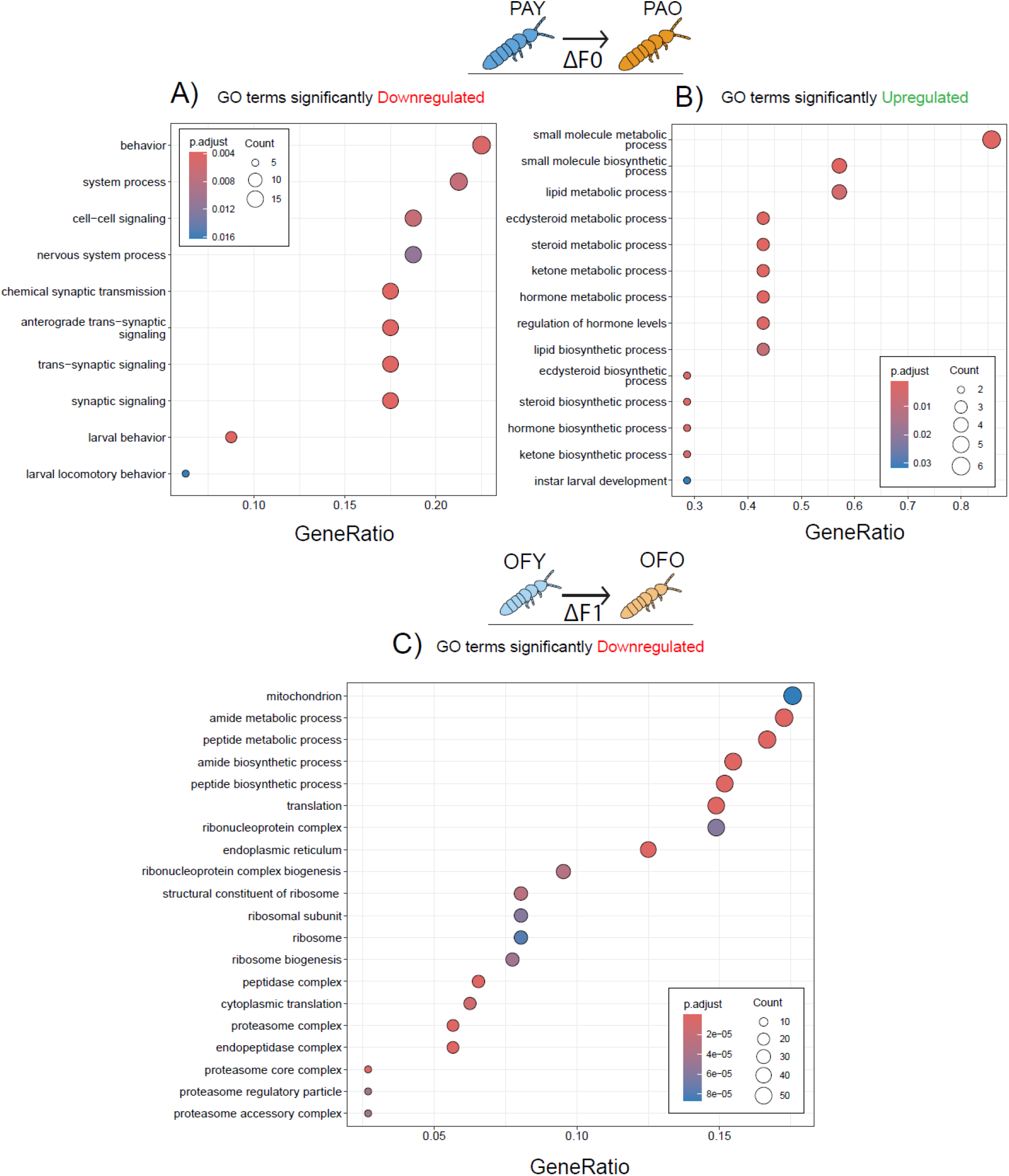
Gene Ontology (GO) enrichment results for differentially expressed genes in older F0 females, and in their F1 offspring at age 20-25 days. Panels A and B show enriched GO terms among genes that were significantly downregulated (A) and upregulated (B) in older mothers relative to young mothers (ΔF0). Panel C shows enriched GO terms for genes significantly downregulated in the offspring of older mothers compared to offspring of young mothers (ΔF1). There were no significantly upregulated GO terms in ΔF1.

We found 17 significantly enriched KEGG pathways in our F0 gene set enrichment analysis (GSEA; adjusted *p* < 0.05; see Figure 4a). Six metabolic and signalling KEGG pathways were downregulated in older *F. candida* females, including the citrate cycle (TCA cycle), oxidative phosphorylation, carbon metabolism, and sphingolipid metabolism. These pathways are necessary for energy production, biosynthesis, and membrane signalling, and their reduced activity suggests a general metabolic decline with age. Additionally, the downregulation of cytoskeletal components and neuroactive ligand-receptor interactions may reflect decreased locomotor activity and reduced sensory responsiveness in older individuals, consistent with the age-related sarcopenia reported in other taxa (Demontis et al. 2013). Eleven KEGG pathways were significantly upregulated in older *F. candida* females, suggesting a broad transcriptional response to age-related physiological demands. The most significantly upregulated KEGG pathways were involved in chromatin organization and regulation of gene expression. These include ATP-dependent chromatin remodelling, the Polycomb repressive complex, and basal transcription factors. The upregulation of these pathways points to a potential reorganization of chromatin architecture and regulation of transcription as individuals age. Additionally, pathways involved in genomic maintenance such as nucleotide excision repair, base excision repair, homologous recombination, the Fanconi anemia pathway, and DNA replication were upregulated, indicating potentially increased investment in DNA damage monitoring and repair mechanisms. Increased activity in the ubiquitin-mediated proteolysis pathway also suggests that older females have a greater need for breaking down proteins, potentially due to accumulation of damaged or misfolded proteins. Two metabolic pathways, ether lipid metabolism and alpha-linolenic acid metabolism, were also upregulated, suggesting shifts in lipid processing and membrane composition in aging tissues. This shift away from carbohydrate-based energy metabolism towards lipid and ketone utilisation is consistent with metabolic reprogramming observed in ageing mammals, where reduced glycolytic and TCA cycle activity is offset by increased reliance on alternative energy substrates (Victoria et al. 1981; Mutlu et al. 2021). Such a shift may reflect compensatory mechanisms to maintain energy homeostasis under conditions of mitochondrial inefficiency. These findings show a multifaceted response in aging *F. candida* that may include increased investment in DNA repair, chromatin remodelling, proteostasis, and metabolism, potentially to counteract age-associated cellular stress.

**Figure 4.**
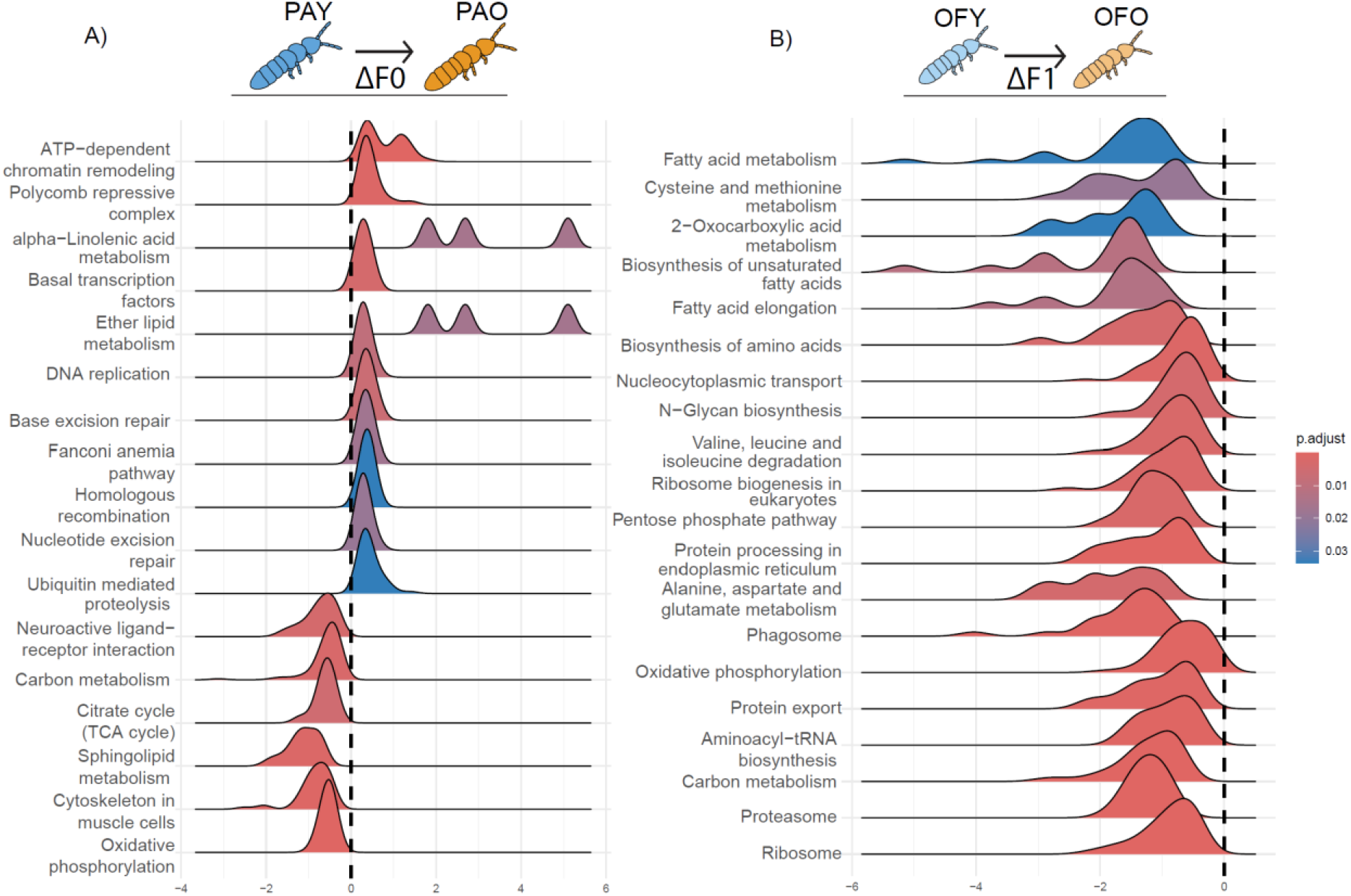
Ridgeplot of gene set enrichment analysis (GSEA) results in the parental (A) and offspring (B) generations. The X-axis represents log_2_ fold change (log_2_FC) values, while the Y-axis lists significantly enriched KEGG pathways. Each ridge reflects the distribution of log_2_FC values for genes within a given pathway, highlighting the direction and strength of enrichment.

### Maternal senescence reprograms offspring gene expression

Maternal (F0) age at reproduction affected gene expression in offspring (F1). Among offspring, a total of 3265 genes were identified as DEGs (ΔF1) in older mothers’ offspring (OFO) compared with young mothers’ offspring (OFY). Of these, 45.2% (1477) were upregulated and 54.8% (1788) were downregulated in OFO relative to OFY (See Figure 2d). Because all offspring were sampled as young adults (age 20–25 days), this transcriptional plasticity suggests considerable developmental sensitivity to transmitted epigenetic states. The homogeneity of group dispersions did not differ between offspring of older mothers (OFO) versus offspring of young mothers (OFY), but the centroids of transcriptional profiles differed significantly between OFY and OFO (PERMANOVA, p < 0.005; see Figure 2b and Supplementary Information).

Of the 3265 differentially expressed genes in offspring, 14.4% (473 genes,150 upregulated, 323 downregulated) were able to be annotated with one-to-one orthologs in Arthropoda, and had GO terms assigned. GO terms significantly downregulated in the offspring of older females were predominantly associated with protein synthesis, processing, and transport. These included processes such as translation, peptide and amide metabolism/biosynthesis, ribosomal structural proteins, endoplasmic reticulum (ER), and components of the proteasome complex. Notably, genes associated with mitochondrial function were also significantly downregulated in the OFO group. These findings suggest an overall reduction in proteostatic activity and a potential decline in mitochondrial function in the offspring of aged adults. No GO terms showed significant upregulation in OFO versus OFY.

KEGG pathway gene set enrichment analysis for the offspring (F1) also revealed a broad downregulation of metabolic and biosynthetic pathways in OFO compared to OFY, with no pathways showing significant upregulation (see Figure 4b). Notably, several core cellular processes were downregulated, including ribosome and ribosome biogenesis, proteasome, oxidative phosphorylation, and carbon metabolism. Pathways involved in amino acid biosynthesis and degradation (e.g., valine, leucine and isoleucine degradation, biosynthesis of amino acids, cysteine and methionine metabolism), fatty acid metabolism (including biosynthesis of unsaturated fatty acids, fatty acid elongation, and fatty acid metabolism), as well as nucleocytoplasmic transport, protein processing and export, and N-glycan biosynthesis, were also significantly downregulated (adjusted *p* < 0.05; Figure 4b). These results suggest a general reduction in biosynthetic capacity and cellular metabolic activity in the offspring of older mothers. Notably, the oxidative phosphorylation and carbon metabolism KEGG pathways were significantly downregulated in both the offspring (F1) and parental (F0) generations, indicating that age-related declines in energy production and central metabolic pathways may be transmitted across generations.

GO and KEGG analyses based on eukaryote-wide annotations yielded broadly similar results, but additionally suggested downregulation of the TOR pathway in offspring of older adults (see Supplementary Information, Table S1, Fig. S1).

### Concordance across generations

To investigate whether offspring of older mothers exhibit gene expression patterns that are concordant with the senescent gene expression of their mothers (ΔF1 ∼ ΔF0), we calculated Pearson’s correlation coefficients (*r*) between ΔF1and ΔF0 based on log_2_ fold changes (log_2_FC) for differentially expressed genes and all genes (Fig. 5a, b). Across all 22,555 genes, we found a significant positive correlation between offspring and parental log_2_FC values (*r* = 0.14, p<0.001; Figure 5b), indicating that age-related changes in maternal gene expression tend to be mirrored from a young age across the offspring transcriptome. Much stronger concordance was observed for the subset of genes showing significant fold-changes in both maternal and offspring generations, including genes upregulated in both generations (*r* = 0.90, p<0.001) and genes downregulated in both generations (*r* = 0.37, p<0.001). These results show that older females transmit a broad syndrome of senescence-like gene expression patterns to their offspring. Notably, the subset of genes that were significantly downregulated in older mothers but upregulated in their offspring showed negative correlation of fold-changes, suggesting compensatory mechanisms that may counteract some age-related effects across generations (*r* = -0.55, p<0.001; See Table S3). No correlation of fold-changes was observed for genes that were upregulated in older mothers but downregulated in their offspring (*r* = 0.03, p > 0.95; Fig. 5b). Simulation analyses confirmed that these cross-generational expression patterns are highly unlikely to have arisen by chance (Supplementary Information, Figure S2).

**Figure 5.**
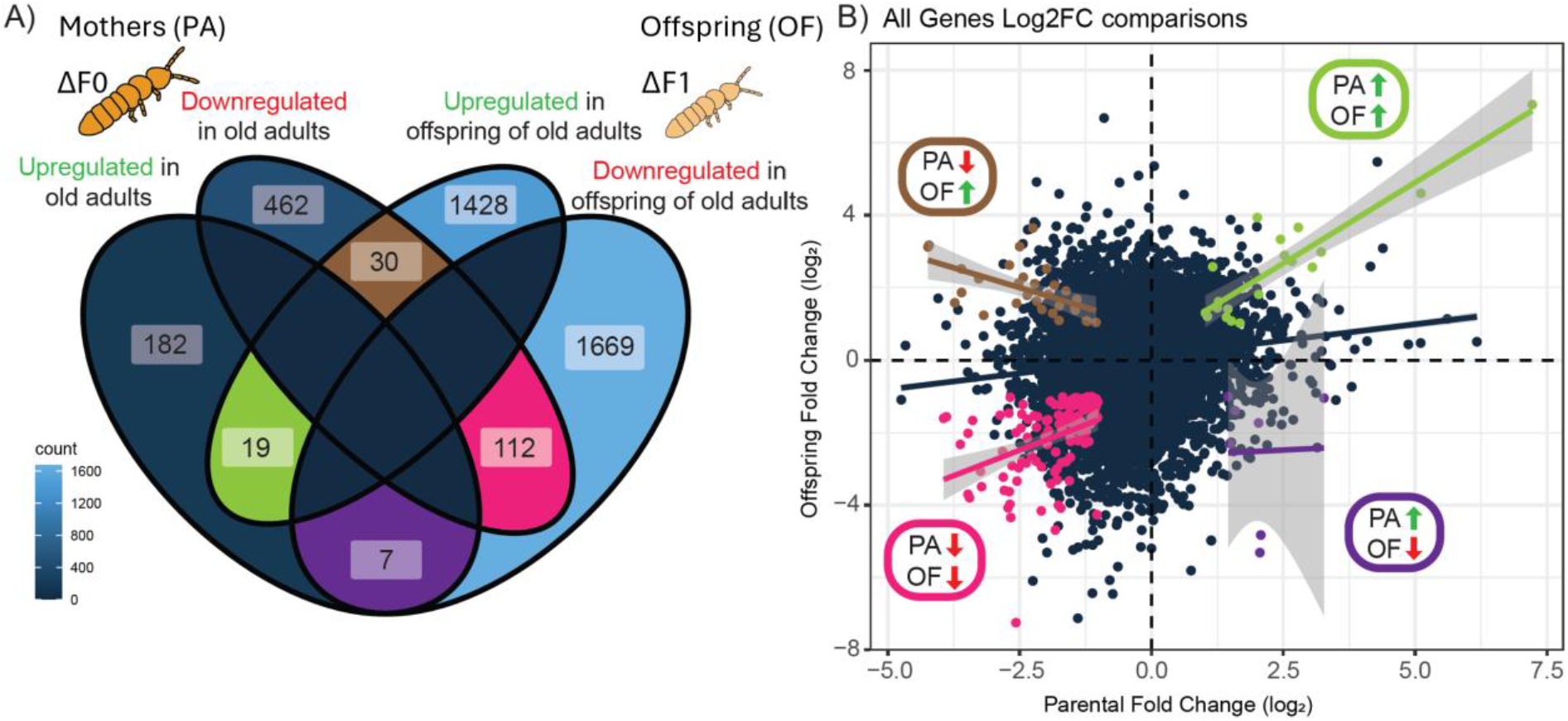
Concordance of fold-change between the parental (F0) generation (ΔF0) and offspring (F1) generation (ΔF1). The Venn diagram (A) shows the number of differentially expressed genes (DEGs) that are unique to each generation or shared between them. The scatterplot (B) displays the log_2_ fold change (log_2_FC) of all genes in the parental (F0) generation (x-axis) versus the offspring (F1) generation (y-axis), with points coloured by gene set: all genes (dark blue), genes upregulated in both generations (green), genes downregulated in both generations (pink), genes upregulated in parents and downregulated in offspring (purple), and genes downregulated in parents and upregulated in offspring (brown).

### Maternal senescence reduces offspring fitness

Maternal age at reproduction had no effect on egg development time (medians: OFY = OFO = 7 d, estimate = 0.068, se = 0.313, *p* = 0.83; Table S5; Figure 6a). However, after hatching, offspring of older mothers took 42% longer to start laying eggs than did offspring of young mothers (medians: OFY = 12 d, OFO = 17 d; estimate = -0.195, se = 0.041, *p* < 0.001; Table S6; Figure 6b).

**Figure 6.**
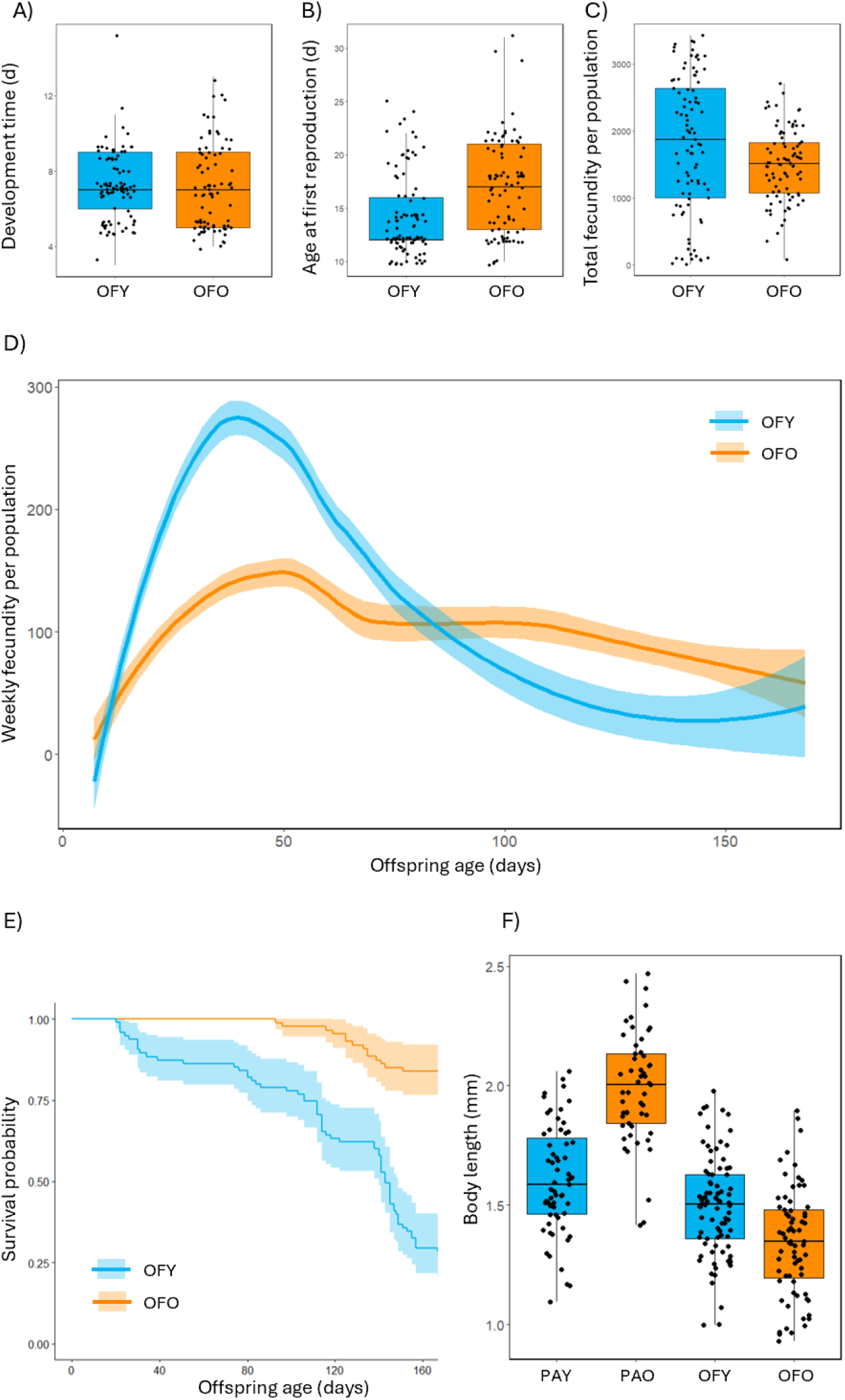
Effects of maternal age at breeding on offspring development time (A), age at first reproduction (B), lifetime (up to 167 days) fecundity (C), age-specific fecundity (D), mortality rate (E) and body size (F). In plots A-C, each point represents one replicate population. In plot F, each point represents one individual. In plot D, loess curves (with standard error bands) are plotted separately for each maternal age treatment. One outlier point was omitted from plots B and C. Plot E shows Kaplan-Meier curves (with standard error bands) for young females’ offspring (OFY) and older females’ offspring (OFO). Plot F shows body size of F0 mothers at young (PAY) and older (PAO) ages, and the body sizes of offspring of young (OFY) and older (OFO) mothers.

Offspring of older mothers laid 16% fewer eggs over their lifetime (up to 167 days) than did offspring of young mothers (means: OFY = 358 eggs, OFO = 300 eggs; estimate = 0.364, se = 0.095, *p* < 0.001; Table S7; Figure 6c), even though the reproductive lifespan of offspring of older mothers was 11% longer than that of offspring of young mothers (medians: OFY = 122 days, OFO = 135 days). Young mothers’ offspring thus achieved a mean daily reproductive rate 55% higher than that of older mothers’ offspring (estimate = 0.588, se = 0.061, *p* < 0.001; Table S8). Older mothers’ offspring had lower fecundity than young mothers’ offspring in early life, but this pattern was partially reversed in later life (maternal age × offspring age interaction: Chi-sq = 11.79, df = 1, *p* < 0.001; Table S9, S10; Figure 6d). Older mothers’ offspring had a lower mortality rate than did young mothers’ offspring (Wald test: *W* = 54.08, df = 1, p < 0.001; Table S11; Figure 6e). Older mothers were larger than young mothers (estimate = -0.375, se = 0.037, *p* < 0.001), but older mothers’ offspring were smaller than young mothers’ offspring (estimate = 0.154, se = 0.043, *p* < 0.001; Table S12, S13; Figure 6F). Overall, these results indicate that maternal senescence had a highly deleterious effect on offspring fitness.

## Discussion

Our results reveal that maternal senescence causes extensive reprograming of offspring transcriptional profiles in *F. candida*. Senescent changes in mothers’ gene expression reflected a multifaceted ageing signature, including reduced neural activity, widespread metabolic reprogramming, and increased transcriptional investment in DNA repair, chromatin organization, and proteostasis. In the offspring of older mothers, gene expression was broadly downregulated for processes related to protein synthesis and proteostasis, mitochondrial function and core metabolic pathways, indicating reduced cellular biosynthetic and energetic activity. Intriguingly, four times as many genes showed significant differential expression in offspring of older mothers than in older mothers themselves, highlighting the importance of maternal age in shaping transcriptional profiles in offspring.

Older mothers and their offspring exhibited concordant patterns of differential gene expression across the entire transcriptome, and especially for loci showing significant differential expression of similar sign in both generations. For loci of known function, observed fold-changes pointed to reduced capacity for energy production and metabolism in both older mothers and in their offspring. Older mothers thus appear to induce a senescence-like transcriptomic syndrome in their offspring via locus-specific transmission of gene expression patterns. Reprogramming of modifiers that drive consistent gene expression changes at multiple loci could also contribute to concordance. Nonetheless, the overlap in genes and pathways exhibiting significant differential expression in maternal and offspring generations was not perfect. In particular, we found significantly downregulated expression in pathways associated with neuronal and muscle function in older mothers but not in their offspring. Moreover, some genes exhibited differential expression of opposite sign in older mothers *versus* in their offspring, and upregulation of some pathways in older mothers was not mirrored in their offspring. Thus, despite broad concordance across generations, some gene expression patterns appear to be unique to older individuals, while other gene expression patterns may be unique to older individuals’ offspring.

Concordant fold-changes between maternal and offspring generations are consistent with germ-line inheritance of a senescent epigenome despite epigenetic reprogramming between generations. This inheritance could involve transmission of senescent patterns of DNA methylation (Hua et al. 2022), histone modification (Wu et al. 2022), noncoding RNA (Li and Casanueva 2016; Zhou et al. 2025), or factors engendering self-sustaining metabolic “loops” (Jablonka and Raz 2009). It remains unknown whether offspring of older mothers not only begin life with a senescence-like gene expression profile but also undergo more rapid changes in gene expression as they age, perhaps as a result of an accelerated “epigenetic clock” (Horvath and Raj 2018).

Maternal senescence was associated with strongly reduced offspring fitness. Offspring of older mothers were smaller, exhibited delayed onset of reproduction, and had lower reproductive rate and lifetime fecundity. These offspring also exhibited longer lifespans, likely associated with their reduced investment in reproduction (Reznick 1985, 1992; Stearns 1998). Our results thus show that maternal senescence can reduce offspring fitness despite extending offspring longevity. Observed effects of maternal senescence on offspring gene expression, such as reduced energy metabolism and protein synthesis, are consistent with the deleterious effects on offspring fitness. The smaller body size of older mothers’ offspring also suggests reduced nutrient provisioning of eggs, which could affect offspring fitness. However, reduced nutrient provisioning is unlikely to explain the senescence-like gene expression patterns that we observed in older mothers’ offspring. There is evidence that deleterious effects of maternal senescence on offspring phenotypes can be amplified by maternal stress (Zaidan et al. 2023) and mitigated by maternal dietary restriction (Gribble et al. 2014; Li et al. 2023). Environment-induced transcriptional changes could mediate such effects.

Transmission of parental senescence to offspring represents a potentially important source of within-population variation in fitness and could have implications for the evolution of ageing (Priest et al. 2002; Bonduriansky and Day 2018; Wylde et al. 2019; Barks and Laird 2020). If older parents produce low-fitness offspring, selection may favour alleles that induce earlier reproduction. However, if offspring of older parents also have extended lifespans, fitness advantages of longer life in environments where background mortality is low could help maintain genetic variation in ageing rates within populations.

In humans and other mammals, older parental age is associated with increased risks of pregnancy failure (Zhang et al. 2022) as well as life-long health problems in offspring (Auroux 1983; Malaspina et al. 2005; Aviv 2012; Wu et al. 2017). Yet, correlational data from human populations also suggest that the effects of parental age are highly multifaceted and complex (Carslake et al. 2017). Correlational studies in human populations and experimental studies on animal models suggest that epigenetic reprogramming/dysregulation contributes to such effects (Teschendorff and Horvath 2025). Our findings provide evidence that such reprogramming involves locus-specific transmission of gene expression across the transcriptome, inducing a senescence-like syndrome from a young age. Research on diverse species is needed to establish how phylogenetically conserved such reprogramming is across animals.

## Acknowledgements

We are grateful to Felipe Floreste for assistance with RNA extraction and processing, and to Thomas Tully for advice on springtail husbandry. We would also like to thank the staff of the Ramaciotti Centre for Genomics. Funding for this research was provided by the Australian Research Council through a Discovery Grant to RB.

## Author contributions

SMM, ZW and RB designed the study. SMM carried out the RNA extractions, conducted the bioinformatic analyses, and wrote the sections of the paper reporting the gene expression analyses. ZW carried out the experiments used to generate samples for gene expression analysis and fitness analysis, processed the fitness data, adapted, tested and deployed the automated image analysis model, and wrote the sections of the paper describing the experiments and fitness assays. RB obtained funding for this research and supervised the project, carried out the fitness analyses, edited the paper.

## References

Amarya, S., K. Singh, and M. Sabharwal. 2018. Ageing Process and Physiological Changes in G. D’Onofrio, A. Greco, and D. Sancarlo, eds. Gerontology. InTech.

Andersen, A. M. N. 2000. Maternal age and fetal loss: Population based register linkage study. BMJ 320:1708–1712.

Ashapkin, V., A. Suvorov, J. R. Pilsner, S. A. Krawetz, and O. Sergeyev. 2023. Age-associated epigenetic changes in mammalian sperm: implications for offspring health and development. Human Reproduction Update 29:24–44.

Auroux, M. 1983. Decrease of learning capacity in offspring with increasing paternal age in the rat. Teratology 27:141–148.

Aviv, A. 2012. Genetics of leukocyte telomere length and its role in atherosclerosis. Mutation Research/Fundamental and Molecular Mechanisms of Mutagenesis 730:68–74.

Barks, P. M. and R. A. Laird. 2020. Parental Age Effects and the Evolution of Senescence. The American Naturalist 195:886–898.

Bartling, B., K. Niemann, R. U. Pliquett, H. Treede, and A. Simm. 2019. Altered gene expression pattern indicates the differential regulation of the immune response system as an important factor in cardiac aging. Experimental Gerontology 117:13–20.

Berchtold, N. C., D. H. Cribbs, P. D. Coleman, J. Rogers, E. Head, R. Kim, T. Beach, C. Miller, J. Troncoso, J. Q. Trojanowski, H. R. Zielke, and C. W. Cotman. 2008. Gene expression changes in the course of normal brain aging are sexually dimorphic. Proceedings of the National Academy of Sciences 105:15605–15610.

Bollati, V., J. Schwartz, R. Wright, A. Litonjua, L. Tarantini, H. Suh, D. Sparrow, P. Vokonas, and A. Baccarelli. 2009. Decline in genomic DNA methylation through aging in a cohort of elderly subjects. Mechanisms of Ageing and Development 130:234–239.

Bonduriansky, R. and C. E. Brassil. 2002. Rapid and costly ageing in wild male flies. Nature 420:377.

Bonduriansky, R. and T. Day. 2018. Extended Heredity: A New Understanding of Inheritance and Evolution. Princeton University Press.

Bongaarts, J., B. S. Mensch, and A. K. Blanc. 2017. Trends in the age at reproductive transitions in the developing world: The role of education. Population Studies 71:139–154.

Bryois, J., A. Buil, P. G. Ferreira, N. I. Panousis, A. A. Brown, A. Viñuela, A. Planchon, D. Bielser, K. Small, T. Spector, and E. T. Dermitzakis. 2017. Time-dependent genetic effects on gene expression implicate aging processes. Genome Research 27:545–552.

Burke, S. N. and C. A. Barnes. 2006. Neural plasticity in the ageing brain. Nature Reviews Neuroscience 7:30–40.

Burkimsher, M. 2015. Europe-wide fertility trends since the 1990s: Turning the corner from declining first birth rates. Demographic Research 32:621–656.

Cannon, L., A. C. Zambon, A. Cammarato, Z. Zhang, G. Vogler, M. Munoz, E. Taylor, J. Cartry, S. I. Bernstein, S. Melov, and R. Bodmer. 2017. Expression patterns of cardiac aging in Drosophila. Aging Cell 16:82–92.

Carlson, K. A., K. Gardner, A. Pashaj, D. J. Carlson, F. Yu, J. D. Eudy, C. Zhang, and L. G. Harshman. 2015. Genome-Wide Gene Expression in relation to Age in Large Laboratory Cohorts of Drosophila melanogaster. Genetics Research International 2015:1–19.

Carslake, D., P. Tynelius, G. Van Den Berg, G. Davey Smith, and F. Rasmussen. 2017. Associations of parental age with health and social factors in adult offspring. Methodological pitfalls and possibilities. Scientific Reports 7:45278.

Choi, S., A. Mc Cartney, D. Park, H. Roberts, T. Brav‐Cubitt, C. Mitchell, and T. R. Buckley. 2024. Multiple hybridization events and repeated evolution of homoeologue expression bias in parthenogenetic, polyploid New Zealand stick insects. Molecular Ecology:e17422.

Chung, H. Y., B. Sung, K. J. Jung, Y. Zou, and B. P. Yu. 2006. The Molecular Inflammatory Process in Aging. Antioxidants & Redox Signaling 8:572–581.

De Magalhães, J. P., J. Curado, and G. M. Church. 2009. Meta-analysis of age-related gene expression profiles identifies common signatures of aging. Bioinformatics 25:875–881.

Demontis, F., R. Piccirillo, A. L. Goldberg, and N. Perrimon. 2013. Mechanisms of skeletal muscle aging: Insights from Drosophila and mammalian models. Disease Models & Mechanisms:dmm.012559.

Dixon, P. 2003. VEGAN, a package of R functions for community ecology. Journal of Vegetation Science 14:927–930.

Dobin, A., C. A. Davis, F. Schlesinger, J. Drenkow, C. Zaleski, S. Jha, P. Batut, M. Chaisson, and T. R. Gingeras. 2013. STAR: Ultrafast universal RNA-seq aligner. Bioinformatics 29:15–21.

Duan, P., B. Li, C. Li, X. Han, Y. Xu, Y. Xing, and W. Yan. 2015. Effects of delayed motherhood on hippocampal gene expression in offspring rats. Molecular and Cellular Biochemistry 405:89–95.

Fountain, M. T. and S. P. Hopkin. 2005. Folsomia candida (Collembola): A “standard” soil arthropod. Annu. Rev. Entomol. 50:201–222.

Fox, J. and S. Weisberg. 2019. An R Companion to Applied Regression. Sage, Thousand Oaks, CA.

Frenk, S. and J. Houseley. 2018. Gene expression hallmarks of cellular ageing. Biogerontology 19:547–566.

Glass, D., A. Viñuela, M. N. Davies, A. Ramasamy, L. Parts, D. Knowles, A. A. Brown, Å. K. Hedman, K. S. Small, A. Buil, E. Grundberg, A. C. Nica, P. Di Meglio, F. O. Nestle, M. Ryten, U. K. B. E. c. the, T. c. the Mu, R. Durbin, M. I. McCarthy, and T. D. Spector. 2013. Gene expression changes with age in skin, adipose tissue, blood and brain. Genome Biology 14:R75.

Gribble, K. E., G. Jarvis, M. Bock, and D. B. M. Welch. 2014. Maternal caloric restriction partially rescues the deleterious effects of advanced maternal age on offspring. Aging Cell 13:623–630.

Hartig, F. 2025. DHARMa: Residual Diagnostics for Hierarchical (Multi-Level / Mixed) Regression Models.

Horvath, S. 2013. DNA methylation age of human tissues and cell types. Genome Biology 14:3156.

Horvath, S. and K. Raj. 2018. DNA methylation-based biomarkers and the epigenetic clock theory of ageing. Nature Reviews Genetics 19:371–384.

Horvath, S., Y. Zhang, P. Langfelder, R. S. Kahn, M. P. Boks, K. Van Eijk, L. H. Van Den Berg, and R. A. Ophoff. 2012. Aging effects on DNA methylation modules in human brain and blood tissue. Genome Biology 13:R97.

Hua, L., W. Chen, Y. Meng, M. Qin, Z. Yan, R. Yang, Q. Liu, Y. Wei, Y. Zhao, L. Yan, and J. Qiao. 2022. The combination of DNA methylome and transcriptome revealed the intergenerational inheritance on the influence of advanced maternal age. Clinical and translational medicine 12:e990.

Hughes, K. A., J. A. Alipaz, J. M. Drnevich, and R. M. Reynolds. 2002. A test of evolutionary theories of aging. Proceedings of the National Academy of Sciences 99:14286–14291.

Ivimey-Cook, E. and J. Moorad. 2020. The diversity of maternal-age effects upon pre-adult survival across animal species. Proceedings of the Royal Society B: Biological Sciences 287:20200972.

Jablonka, E. and G. Raz. 2009. Transgenerational Epigenetic Inheritance: Prevalence, Mechanisms, and Implications for the Study of Heredity and Evolution. The Quarterly Review of Biology 84:131–176.

Janssens, G. E., A. C. Meinema, J. González, J. C. Wolters, A. Schmidt, V. Guryev, R. Bischoff, E. C. Wit, L. M. Veenhoff, and M. Heinemann. 2015. Protein biogenesis machinery is a driver of replicative aging in yeast. eLife 4:e08527.

Jiang, Y., H. Zhang, S. Chen, S. Ewart, J. W. Holloway, H. Arshad, and W. Karmaus. 2024. Intergenerational association of DNA methylation between parents and offspring. Scientific Reports 14:19812.

Jocker, G., A. Chaurasia, and J. Qui. 2023. YOLO by Ultralytics. Journal of Computer and Communications 11:100–110.

Jones, O., A. Scheuerlein, R. Salguero-Gómez, C. Giovanni Camarda, R. Schaible, B. B. Casper, J. P. Dahlgren, J. Ehrlén, M. B. García, E. S. Menges, P. F. Quintana-Ascencio, H. Caswell, A. Baudisch, and J. W. Vaupel. 2014. Diversity of ageing across the tree of life. Nature 505:169–173.

Kamei, Y., Y. Tamada, Y. Nakayama, E. Fukusaki, and Y. Mukai. 2014. Changes in Transcription and Metabolism During the Early Stage of Replicative Cellular Senescence in Budding Yeast. Journal of Biological Chemistry 289:32081–32093.

Kawai, K., T. Harada, T. Ishikawa, R. Sugiyama, T. Kawamura, A. Yoshida, O. Tsutsumi, F. Ishino, T. Kubota, and T. Kohda. 2018. Parental age and gene expression profiles in individual human blastocysts. Scientific Reports 8:2380.

Keefe, D. L., S. Franco, L. Liu, J. Trimarchi, B. Cao, S. Weitzen, S. Agarwal, and M. A. Blasco. 2005. Telomere length predicts embryo fragmentation after in vitro fertilization in women—Toward a telomere theory of reproductive aging in women. American Journal of Obstetrics and Gynecology 192:1256–1260.

Kim, H. J., K. J. Jung, B. P. Yu, C. G. Cho, J. S. Choi, and H. Y. Chung. 2002. Modulation of redox-sensitive transcription factors by calorie restriction during aging. Mechanisms of Ageing and Development 123:1589–1595.

Kumar, A., J. R. Gibbs, A. Beilina, A. Dillman, R. Kumaran, D. Trabzuni, M. Ryten, R. Walker, C. Smith, B. J. Traynor, J. Hardy, A. B. Singleton, and M. R. Cookson. 2013. Age-associated changes in gene expression in human brain and isolated neurons. Neurobiology of Aging 34:1199–1209.

Lansing, A. I. 1947. A transmissible, cumulative, and reversible factor in aging. Journal of Gerontology 2:228–239.

Lansing, A. I. 1954. A nongenetic factor in the longevity of rotifers. Annals of the New York Academy of Sciences 57:455–464.

Lee, J. S., W. O. Ward, H. Ren, B. Vallanat, G. J. Darlington, E. S. Han, J. C. Laguna, J. H. DeFord, J. Papaconstantinou, C. Selman, and J. C. Corton. 2012. Meta-analysis of gene expression in the mouse liver reveals biomarkers associated with inflammation increased early during aging. Mechanisms of Ageing and Development 133:467–478.

Li, C. and O. Casanueva. 2016. Epigenetic inheritance of proteostasis and ageing. Essays in Biochemistry 60:191–202.

Li, C., H. Zhang, H. Wu, R. Li, D. Wen, Y. Tang, Z. Gao, R. Xu, S. Lu, Q. Wei, X. Zhao, M. Pan, and B. Ma. 2023. Intermittent fasting reverses the declining quality of aged oocytes. Free radical biology & medicine 195:74–88.

López-Otín, C., M. A. Blasco, L. Partridge, M. Serrano, and G. Kroemer. 2013. The Hallmarks of Aging. Cell 153:1194–1217.

López-Otín, C., M. A. Blasco, L. Partridge, M. Serrano, and G. Kroemer. 2023. Hallmarks of aging: An expanding universe. Cell 186:243–278.

Lu, Z., Z. Zhang, Z. Xu, A. Abdulraouf, W. Zhou, and J. Cao. 2026. Organism-wide cellular dynamics and epigenomic remodeling in mammalian aging. Science 391:eadw6273.

Lüdecke, D., M. Ben-Shachar, I. Patil, P. Waggoner, and D. Makowski. 2021. performance: An R Package for Assessment, Comparison and Testing of Statistical Models. Journal of Open Source Software 6:3139.

Ma, X., G. Zhan, M. C. Sleumer, S. Chen, W. Liu, M. Q. Zhang, and X. Liu. 2016. Analysis of C. elegans muscle transcriptome using trans-splicing-based RNA tagging (SRT). Nucleic Acids Research:gkw734.

Malaspina, D., A. Reichenberg, M. Weiser, S. Fennig, M. Davidson, S. Harlap, R. Wolitzky, J. Rabinowitz, E. Susser, and H. Y. Knobler. 2005. Paternal age and intelligence: Implications for age-related genomic changes in male germ cells. Psychiatric Genetics 15:117–125.

McGillycuddy, M., D. I. Warton, G. Popovic, and B. M. Bolker. 2025. Parsimoniously Fitting Large Multivariate Random Effects in glmmTMB. Journal of Statistical Software 112.

Medawar, P. B. 1952. An Unsolved problem of biology: an inaugural lecture delivered at university college, London, 6 December, 1951. H.K. Lewis and Company.

Miller, S. M., Z. Wylde, and R. Bonduriansky. 2026. Data and code for Maternal senescence bradly reprograms gene expression in offspring. Zenodo [link to be added after acceptance].

Mirza, N., K. Pollock, D. B. Hoelzinger, A. L. Dominguez, and J. Lustgarten. 2011. Comparative kinetic analyses of gene profiles of naïve CD4+ and CD8+ T cells from young and old animals reveal novel age‐related alterations. Aging Cell 10:853–867.

Monaghan, P., A. A. Maklakov, and N. B. Metcalfe. 2020. Intergenerational Transfer of Ageing: Parental Age and Offspring Lifespan. Trends in Ecology & Evolution 35:927–937.

Mutlu, A. S., J. Duffy, and M. C. Wang. 2021. Lipid metabolism and lipid signals in aging and longevity. Developmental Cell 56:1394–1407.

Ntostis, P., D. Iles, G. Kokkali, T. Vaxevanoglou, E. Kanavakis, A. Pantou, J. Huntriss, K. Pantos, and P. M. 2022. The impact of maternal age on gene expression during the GV to MII transition in euploid human oocytes. Human Reproduction Update 37:80–92.

Oecd. 2016. Test No. 232: Collembolan Reproduction Test in Soil. OECD Guidelines for the Testing of Chemicals, Sectionv2. OECD Publishing, Paris.

Oecd. 2024. Society at a Glance 2024: OECD Social Indicators. OECD.

Pellestor, F., B. Andréo, F. Arnal, C. Humeau, and J. Demaille. 2003. Maternal aging and chromosomal abnormalities: New data drawn from in vitro unfertilized human oocytes. Human Genetics 112:195–203.

Perez, M. F. and B. Lehner. 2019. Intergenerational and transgenerational epigenetic inheritance in animals. Nature Cell Biology 21:143–151.

Peters, M. J., R. Joehanes, L. C. Pilling, C. Schurmann, K. N. Conneely, J. Powell, E. Reinmaa, G. L. Sutphin, A. Zhernakova, K. Schramm, Y. A. Wilson, S. Kobes, T. Tukiainen, N. U. Consortium, M. A. Nalls, D. G. Hernandez, M. R. Cookson, R. J. Gibbs, J. Hardy, and A. D. Johnson. 2015. The transcriptional landscape of age in human peripheral blood. Nature Communications 6:8570.

Philipp, O., A. Hamann, J. Servos, A. Werner, I. Koch, and H. D. Osiewacz. 2013. A Genome-Wide Longitudinal Transcriptome Analysis of the Aging Model Podospora anserine. PLoS ONE 8:e83109.

Priest, N. K., B. Mackowiak, and D. E. L. Promislow. 2002. The Role of Parental Age Effects on the Evolution of Aging. Evolution 56:927–935.

Reznick, D. 1985. Costs of Reproduction: An Evaluation of the Empirical Evidence. Oikos 44:257.

Reznick, D. 1992. Measuring the Costs of Reproduction. Trends in Ecology & Evolution 7:42–45.

Robinson, M. D., D. J. McCarthy, and G. K. Smyth. 2010. edgeR: A Bioconductor package for differential expression analysis of digital gene expression data. Bioinformatics 26:139–140.

Ruhmann, H., M. Koppik, M. F. Wolfner, and C. Fricke. 2018. The impact of ageing on male reproductive success in Drosophila melanogaster. Experimental Gerontology 103:1–10.

Schneider, C. A., W. S. Rasband, and K. W. Eliceiri. 2012. NIH Image to ImageJ: 25 years of image analysis. Nature Methods 9:671–675.

Schumacher, B., I. Van Der Pluijm, M. J. Moorhouse, T. Kosteas, A. R. Robinson, Y. Suh, T. M. Breit, H. Van Steeg, L. J. Niedernhofer, W. Van Ijcken, A. Bartke, S. R. Spindler, J. H. J. Hoeijmakers, G. T. J. Van Der Horst, and G. A. Garinis. 2008. Delayed and Accelerated Aging Share Common Longevity Assurance Mechanisms. PLoS Genetics 4:e1000161.

Serre, V. and B. Robaire. 1998. Paternal age affects fertility and progeny outcome in the Brown Norway rat. Fertility and Sterility 70:625–631.

Stearns, S. C. 1998. Reproductive Lifespan And Ageing. Pp. 180–205. The Evolution Of Life Histories. Oxford University PressOxford.

Steuerwald, N. M., M. G. Bermúdez, D. Wells, S. Munné, and J. Cohen. 2007. Maternal age-related differential global expression profiles observed in human oocytes. Reproductive BioMedicine Online 14:700–708.

Terao, A., A. Apte-Deshpande, L. Dousman, S. Morairty, B. P. Eynon, T. S. Kilduff, and Y. R. Freund. 2002. Immune response gene expression increases in the aging murine hippocampus. Journal of Neuroimmunology 132:99–112.

Teschendorff, A. E. and S. Horvath. 2025. Epigenetic ageing clocks: statistical methods and emerging computational challenges. Nature Reviews Genetics 26:350–368.

Therneau, T. M. 2024. coxme: Mixed Effects Cox Models.

Van Den Akker, E. B., W. M. Passtoors, R. Jansen, E. W. Van Zwet, J. J. Goeman, M. Hulsman, V. Emilsson, M. Perola, G. Willemsen, B. W. J. H. Penninx, B. T. Heijmans, A. B. Maier, D. I. Boomsma, J. N. Kok, P. E. Slagboom, M. J. T. Reinders, and M. Beekman. 2014. Meta‐analysis on blood transcriptomic studies identifies consistently coexpressed protein– protein interaction modules as robust markers of human aging. Aging Cell 13:216–225.

Victoria, J., J. Satrústegui, and A. Machado. 1981. Metabolic Implications of Ageing: Changes in Activities of KeyLipogenic and Gluconeogenic Enzymes in the Aged Rat Liver. Enzyme 26:144–152.

Webb, M. and D. P. Sideris. 2020. Intimate Relations—Mitochondria and Ageing. International Journal of Molecular Sciences 21:7580.

Williams, C. R., A. Baccarella, J. Z. Parrish, and C. C. Kim. 2016. Trimming of sequence reads alters RNA-Seq gene expression estimates. BMC Bioinformatics 17:103.

Wu, S., F. Wu, Y. Ding, J. Hou, J. Bi, and Z. Zhang. 2017. Advanced parental age and autism risk in children: A systematic review and meta‐analysis. Acta Psychiatrica Scandinavica 135:29–41.

Wu, Y.-W., S. Li, W. Zheng, Y.-C. Li, L. Chen, Y. Zhou, Z.-Q. Deng, G. Lin, H.-Y. Fan, and Q.-Q. Sha. 2022. Dynamic mRNA degradome analyses indicate a role of histone H3K4 trimethylation in association with meiosis-coupled mRNA decay in oocyte aging. Nature Communications 13:3191.

Wylde, Z., F. Spagopoulou, A. K. Hooper, A. A. Maklakov, and R. Bonduriansky. 2019. Parental breeding age effects on descendants’ longevity interact over two generations in matrilines and patrilines. PLoS Biology 17:e3000556.

Yeung, E., R. J. Biedrzycki, L. C. Gómez Herrera, P. Issarapu, J. Dou, I. F. Marques, S. R. Mansuri, C. M. Page, J. Harbs, D. Khodasevich, E. Poisel, Z. Niu, C. Allard, E. Casey, F. M. Berstein, G. Mancano, H. R. Elliott, R. Richmond, Y. He, J. Ronkainen, S. Sebert, E. M. Bell, G. Sharp, S. L. Mumford, E. F. Schisterman, G. R. Chandak, C. H. D. Fall, S. A. Sahariah, M. J. Silver, A. M. Prentice, L. Bouchard, M. Domellof, C. West, N. Holland, A. Cardenas, B. Eskenazi, L. Zillich, S. H. Witt, T. Send, Breton. C, K. M. Bakulski, M. D. Fallin, R. J. Schmidt, D. J. Stein, H. J. Zar, V. W. V. Jaddoe, J. Wright, R. Grazuleviciene, K. B. Gutzkow, J. Sunyer, A. Huels, M. Vrijheid, S. Harlid, S. London, M. F. Hivert, J. Felix, M. Bustamante, and W. Guan. 2024. Maternal age is related to offspring DNA methylation: A meta-analysis of results from the PACE consortium. Aging Cell 8:e14194.

Yoshizaki, K., R. Kimura, H. Kobayashi, S. Oki, T. Kikkawa, L. Mai, K. Koike, K. Mochizuki, H. Inada, Y. Matsui, T. Kono, and N. Osumi. 2021. Paternal age affects offspring via an epigenetic mechanism involving REST/NRSF. The EMBO Reports 22:EMBR202051524.

Zahn, J. M., R. Sonu, H. Vogel, E. Crane, K. Mazan-Mamczarz, R. Rabkin, R. W. Davis, K. Becker, A. B. Owen, and S. K. Kim. 2005. Transcriptional profiling of aging in human muscle reveals a common aging signature. PLoS Genetics:e115.

Zaidan, H., A. Wnuk, I. M. Aderka, M. Kajta, and I. Gaisler-Salomon. 2023. Pre-reproductive stress in adolescent female rats alters maternal care and DNA methylation patterns across generations. Stress 26:2201325.

Zhang, C., L. Yan, and J. Qiao. 2022. Effect of advanced parental age on pregnancy outcome and offspring health. Journal of Assisted Reproduction and Genetics 39:1969–1986.

Zhou, Q., G. Li, H. Ji, J. Ji, Z. Ling, J. Sun, H. Ding, J. Zhang, X. Ling, X. Chen, and L. Xu. 2025. Age-related changes in lncRNA expression in sperm. Translational andrology and urology 14:1408–1417.

