## Supplementary information, tables and figures for "Maternal senescence broadly reprograms gene expression in offspring"

### GO analysis based on eukaryote-wide annotations

DEGs were functionally annotated with Gene Ontology (GO) terms using the orthology-based annotation tool eggNOG-mapper (emapper v2.1.12; http://eggnog-mapper.embl.de) with default settings. In the main analysis, we retained only annotations with a maximum taxonomic specificity within Arthropoda, which resulted in the exclusion of a substantial number of potentially informative annotations. We therefore additionally performed GO analyses using the full set of eukaryotic annotations, excluding only annotations with maximum taxonomic specificity restricted to bacteria, fungi, or viruses (i.e., annotations unique to these clades and not conserved across broader lineages). Of the 812 differentially expressed genes in ΔF0, 200 (24.6%) were able to be annotated with GO terms with this broader database.

Four GO terms, steroid metabolic process, ketone metabolic process, hormone metabolic process, and ecdysteroid metabolic process, were significantly upregulated in ΔF0 (adjusted $p < 0.05$; see Fig. S1b). This was broadly consistent with arthropod-restricted annotations reported in the main text, supporting an age-associated shift in metabolic activity in adult individuals. In contrast, 28 GO terms were significantly enriched among downregulated genes in ΔF0 (adjusted $p < 0.05$), which again broadly aligned with the arthropod-based annotation analysis, particularly with respect to reduced synaptic signalling. However, the eukaryote-wide annotation set also included significantly enriched GO terms associated with the development of anatomical structures not present in *Folsomia candida*, such as lung, kidney, and inner ear morphogenesis (see Fig. S1a). The function of these pathways in *F. candida* is not clear.

Of the total of 3265 differentially expressed genes in ΔF1, 1103 (33.7%) were able to be annotated with GO terms. GO terms significantly downregulated in the offspring of old adults were predominantly associated with protein synthesis, processing, and transport, consistent with the arthropod-restricted annotations reported in the main text. In contrast, upregulated GO terms in ΔF1 were enriched for pathways involved in metabolism, immune response, and development, a pattern not seen in the arthropod-restricted annotation GO analysis. Metabolic processes included steroid metabolic process, TOR and TORC1 signalling, and cellular responses to insulin and VEGF stimuli, indicating altered nutrient sensing and hormonal signalling. Additionally, genes associated with regulation of eating behaviours were significantly upregulated. Immune-related terms such as hemocyte development and proliferation and monocyte chemotaxis were also enriched, suggesting altered immune activity. Developmental processes, including genital disc morphogenesis, Malpighian tubule development, and eye morphogenesis, were also significantly upregulated, pointing to potential changes in developmental trajectories. Of the specific genes upregulated within the TORC signalling pathways, many were identified as inhibitors of TORC1 activity (see Table S1).

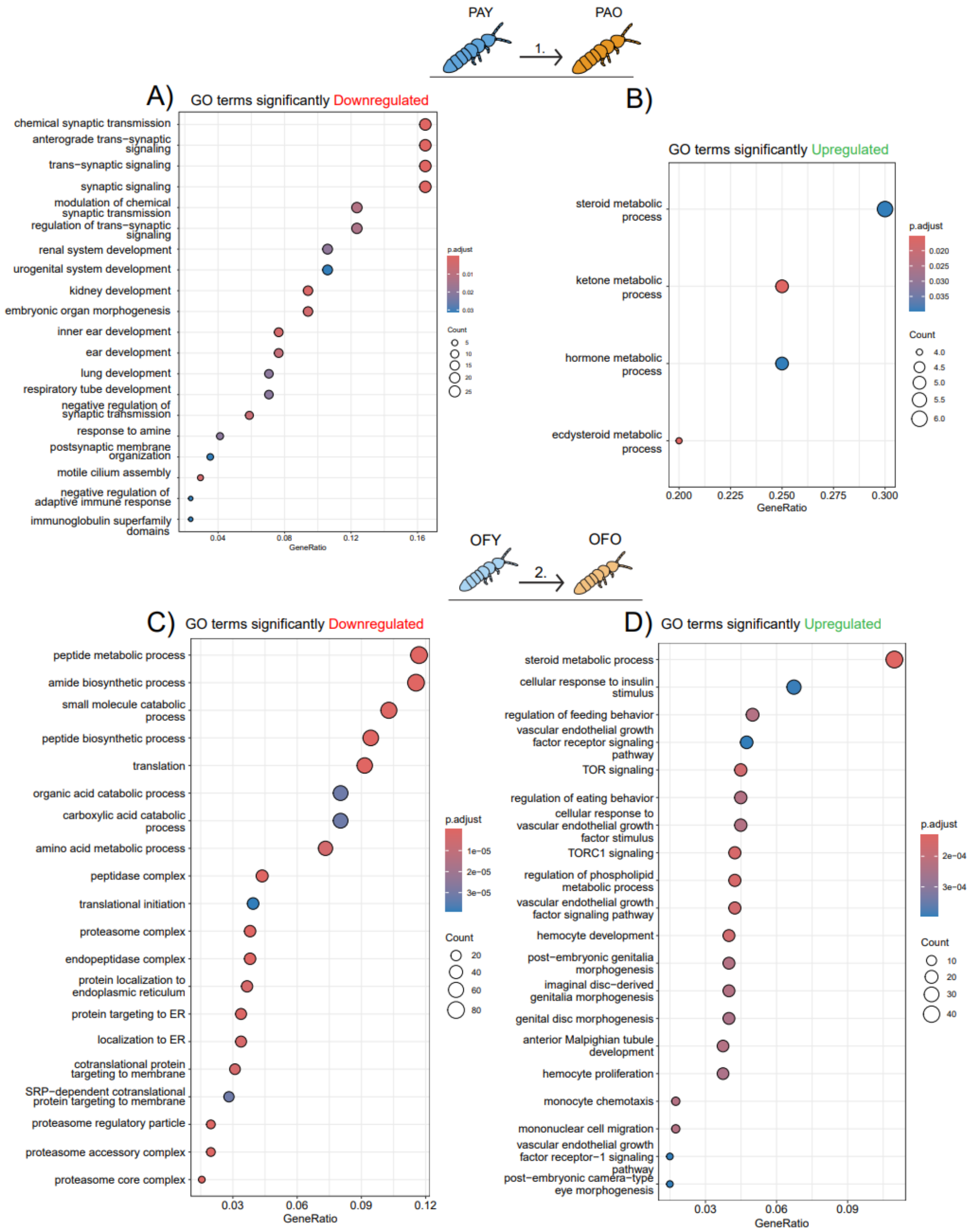


**Figure S1:** Gene Ontology (GO) enrichment results for differentially expressed genes in ΔF0, and in ΔF1 based on eukaryote-wide annotations. Panels A and B show enriched Gene Ontology (GO) terms among genes that were significantly downregulated (A) and upregulated (B) in aged mothers relative to young mothers. Panels C and D show enriched GO terms for genes downregulated (C) and upregulated (D) in the offspring of aged mothers compared to offspring of young mothers. Note that some of the enriched GO terms relate to vertebrate organ systems that are lacking in *F. candida*. The functions of the orthologous loci in *F. candida* are not known.

**Table S1:** Genes associated with TORC signalling that were significantly upregulated in offspring of older mothers compared to offspring of young mothers, based on eukaryote-wide annotations. Log2FC refers to the log2 fold change in the ΔF1 comparison.

| **Gene in *Folsomia candida*** | **Ortholog gene name** | **Log2FC** | **Function** | **Citation** |
|---|---|---|---|---|
| LOC110847979 | IRS1 | 1.193 | Is a central adapter protein in the insulin signalling pathway and is an **upstream activator** and feedback-regulated target of TORC signalling. | Destefano and Jacinto 2013[1] |
| LOC110842942 | SIRT1 | 1.198 | Is a NAD+ dependent deacetylase. It is a **negative** regulator of mTORC signalling. | Ghosh McBurney, and Robbins 2010[2] |
| LOC110852443 | PRKAG2 | 1.21 | PRKAG2 encodes the gamma-2 subunit of AMP-activated protein kinase (AMPK). It is directly involved in the mTOR signalling pathway as AMPK activation is known to inhibit mTORC1 activity. It is typically acting as a **negative regulator** of mTORC1. | Smiles et al 2024[3] |
| LOC118436879 | EIF2AK2 | 1.90 | Encodes the protein kinase PKR (Protein Kinase R), one of the four main kinases that phosphorylate eIF2α in response to cellular stress (e.g. oxidative stress). eIF2α phosphorylation **inhibits** general translation as a protective response to stress. TORC and PKR both regulate protein synthesis but TORC promotes it while PKR represses it. | Wek 2018[4] Proud 2009[5] |
| LOC110848463 | RPTOR | 1.09 | RAPTOR is a core component of the mTORC1 complex. | Wang et al 2009[6] |
| LOC118437787 | MAPK1 | 1.43 | MAPK1 is a central kinase in MAPK/ERK signalling. MAPK1 phosphorylates and inhibits TSC2 (Tuberous Sclerosis Complex 2), which suppresses an activator of mTORC1. MAPK1 activation thus leads to **increased** mTORC1 activity. | Brittany L Dunkerly-Eyring et al 2022[7] Kim, Cook, and Chen 2017[8] |

| | | | | |
|---|---|---|---|---|
| LOC110858510 | DDiT4L | 1.32 | DDIT4L is a well-established regulator of the TORC pathway, it **inhibits** TORC1 signalling and promotes autophagy. It can however stimulate TORC2 signalling. | Simonson et al 2017[9] |
| LOC110845943 | ATG9A | 1.61 | ATG9A is a **negative regulator** of TORC. | Wen et al 2017[10] |

**Table S2:** Each RNA-seq sample used in this study, including sample ID, experimental group, isoline, replicate number, total number of reads, number of uniquely mapped reads, and the percentage of uniquely mapped reads to the *Folsomia candida* reference genome (GenBank accession: GCF_002217175.1).

| Sample | Group | Isoline | Replicate | Total number of reads | Uniquely mapped reads | Uniq. Mapping percent |
|---|---|---|---|---|---|---|
| OFO1 | OFO | I1 | 1 | 37469191 | 26932607 | 71.88% |
| OFY1 | OFY | I1 | 1 | 52219801 | 40134606 | 76.86% |
| OFY2 | OFY | I1 | 2 | 40383047 | 30215883 | 74.82% |
| PAO2 | PAO | I1 | 2 | 57756289 | 49387172 | 85.51% |
| PAO1 | PAO | I1 | 1 | 46807029 | 38355002 | 81.94% |
| PAY1 | PAY | I1 | 1 | 44909166 | 38049485 | 84.73% |
| PAY3 | PAY | I1 | 2 | 38417407 | 29132845 | 75.83% |
| OFO6 | OFO | I2 | 2 | 50138737 | 40325585 | 80.43% |
| OFO4 | OFO | I2 | 1 | 37688276 | 27898089 | 74.02% |
| OFY4 | OFY | I2 | 1 | 41101075 | 31097211 | 75.66% |
| OFY5 | OFY | I2 | 2 | 41964254 | 32853539 | 78.29% |
| PAO6 | PAO | I2 | 2 | 53342819 | 44216676 | 82.89% |
| PAO5 | PAO | I2 | 1 | 54106295 | 44185328 | 81.66% |
| PAY5 | PAY | I2 | 1 | 49128608 | 41716813 | 84.91% |
| OFO11 | OFO | I4 | 2 | 44107954 | 32419212 | 73.50% |
| OFO10 | OFO | I4 | 1 | 41316956 | 30530711 | 73.89% |
| OFY12 | OFY | I4 | 2 | 59817326 | 49633140 | 82.97% |
| OFY10 | OFY | I4 | 1 | 38942572 | 30027520 | 77.11% |
| PAO12 | PAO | I4 | 2 | 48115182 | 37444299 | 77.82% |
| PAO11 | PAO | I4 | 1 | 27400117 | 21741513 | 79.35% |
| PAY11 | PAY | I4 | 1 | 33625244 | 27119279 | 80.65% |
| PAY12 | PAY | I4 | 2 | 59588778 | 51196472 | 85.92% |
| OFO15 | OFO | I5 | 1 | 46892514 | 33035104 | 70.45% |
| OFY14 | OFY | I5 | 1 | 32600656 | 23256319 | 71.34% |
| PAO13 | PAO | I5 | 1 | 40756634 | 33019846 | 81.02% |
| PAO14 | PAO | I5 | 2 | 51275128 | 41638243 | 81.21% |
| PAY13 | PAY | I5 | 1 | 53683833 | 45051582 | 83.92% |
| OFO16 | OFO | I6 | 1 | 41611641 | 34525038 | 82.97% |
| OFO17 | OFO | I6 | 2 | 43964084 | 35159223 | 79.97% |
| OFY16 | OFY | I6 | 1 | 21638252 | 16402602 | 75.80% |
| OFY18 | OFY | I6 | 2 | 48224949 | 39620797 | 82.16% |
| PAO16 | PAO | I6 | 1 | 32878335 | 25503443 | 77.57% |
| PAO18 | PAO | I6 | 2 | 52549883 | 44283032 | 84.27% |
| PAY17 | PAY | I6 | 1 | 42449038 | 34630536 | 81.58% |
| OFO21 | OFO | I7 | 2 | 39624767 | 30285050 | 76.43% |
| OFO20 | OFO | I7 | 1 | 39397390 | 28441153 | 72.19% |
| OFY19 | OFY | I7 | 1 | 36770960 | 29901082 | 81.32% |
| OFY20 | OFY | I7 | 2 | 39189908 | 31640087 | 80.74% |

| | | | | | | |
|---|---|---|---|---|---|---|
| PAO20 | PAO | I7 | 2 | 69903435 | 60058296 | 85.92% |
| PAO19 | PAO | I7 | 1 | 47646944 | 39549279 | 83.00% |
| PAY19 | PAY | I7 | 1 | 55449009 | 45960396 | 82.89% |
| PAY20 | PAY | I7 | 2 | 35273387 | 27858883 | 78.98% |
| OFO23 | OFO | I8 | 2 | 34710146 | 26007804 | 74.93% |
| OFO22 | OFO | I8 | 1 | 43906668 | 33518936 | 76.34% |
| OFY22 | OFY | I8 | 1 | 49185225 | 38407742 | 78.09% |
| OFY23 | OFY | I8 | 2 | 43488963 | 34878479 | 80.20% |
| PAO23 | PAO | I8 | 1 | 41532413 | 33850738 | 81.50% |
| PAO24 | PAO | I8 | 2 | 44723460 | 36972331 | 82.67% |
| PAY23 | PAY | I8 | 2 | 50735769 | 41227642 | 81.26% |
| PAY22 | PAY | I8 | 1 | 47720119 | 40494591 | 84.86% |
| OFO25 | OFO | I9 | 1 | 48067245 | 35324051 | 73.49% |
| OFY25 | OFY | I9 | 1 | 40069328 | 30597353 | 76.36% |
| OFY27 | OFY | I9 | 2 | 36879407 | 28981533 | 78.58% |
| PAO27 | PAO | I9 | 2 | 43184213 | 36141936 | 83.69% |
| PAO26 | PAO | I9 | 1 | 38173379 | 31104130 | 81.48% |
| PAY27 | PAY | I9 | 2 | 50261740 | 41971801 | 83.51% |
| PAY25 | PAY | I9 | 1 | 41800641 | 34022497 | 81.39% |
| OFO28 | OFO | I10 | 1 | 47256085 | 34779795 | 73.60% |
| OFY29 | OFY | I10 | 2 | 38175736 | 31324608 | 82.05% |
| OFY28 | OFY | I10 | 1 | 46128304 | 38996291 | 84.54% |
| PAO28 | PAO | I10 | 1 | 41061361 | 32938264 | 80.22% |
| PAO29 | PAO | I10 | 2 | 43792805 | 34915293 | 79.73% |
| PAY29 | PAY | I10 | 2 | 46193774 | 37125140 | 80.37% |
| PAY28 | PAY | I10 | 1 | 42336165 | 34365824 | 81.17% |

**Table S3:** Pearson's correlation coefficients and correlation test results comparing $\log_2$ fold changes ($\log_2 FC$) between older versus young females (ΔF0) and offspring of older and young females (ΔF1). Correlations were calculated for all non-differentially expressed genes (n = 22,387), as well as for subsets of genes that were upregulated in both parental (F0) and offspring (F1) generations, downregulated in both generations, or showed opposite regulation between generations. The table reports the number of genes in each group, Pearson's r, test statistic, degrees of freedom (df), p-value, and 95% confidence intervals.

| Group | Number of genes | Pearsons's correlation coefficient | Test statistic | DF | p-value | 95% confidence interval |
|---|---|---|---|---|---|---|
| All genes | 22387 | 0.12 | 18.39 | 22385 | 2.2e-16 | 0.11, 0.14 |
| Upregulated in both F0 and F1 | 19 | 0.90 | 8.65 | 17 | 1.24e-07 | 0.76, 0.96 |
| Downregulated in both F0 and F1 | 112 | 0.37 | 4.23 | 110 | 4.78e-05 | 0.20, 0.52 |
| Upregulated in F0, Downregulated in F1 | 7 | 0.03 | 0.06 | 5 | 0.95 | -0.74, 0.76 |
| Downregulated in F0, Upregulated in F1 | 30 | -0.55 | -3.52 | 28 | 0.001 | -0.76, -0.24 |

**Simulation analysis:**

After identifying genes that were consistently upregulated or downregulated in both parental and offspring comparisons, as well as genes showing opposite regulation between generations, a randomization test was performed to generate a null distribution. Simulations showed that, by chance, an average of approximately 8.5 genes were classified as uniquely differentially expressed as upregulated or downregulated in either generation (See Table S4). This contrasts sharply with the observed counts of 182, 462, 1428, and 1669 genes, demonstrating that these values are highly unlikely to have occurred randomly. Furthermore, the number of genes exhibiting consistent regulation across both generations (or opposite regulation) was, on average, less than one gene in the simulated data (See Table S4; Fig. S1), contrasting with the observed counts of 19, 112, 7, and 30. These results provide evidence for a transcriptomic signature of senescence in the offspring of aged mothers, reflecting intergenerational molecular consequences of maternal age in offspring.

**Table S4:** Comparison of observed and simulated numbers of differentially expressed genes (DEGs) across generations and directions of change. This table shows the number of DEGs identified in the observed dataset alongside the mean number identified in 10,000 iterations of simulated data using a negative binomial model. Comparisons are grouped by the generation(s) in which differential expression occurred and the direction of change (upregulated, downregulated, or opposite between generations). Standard deviation (sd) from the simulations is shown in parentheses.

| Generation(s) | Direction of change | DEGs in observed data | Mean DEGs in simulated data |
|---|---|---|---|
| Parental only | Upregulated | 182 | 8.48 (4.17) |
| Parental only | Downregulated | 462 | 8.49 (4.15) |
| Offspring only | Upregulated | 1428 | 8.45 (4.18) |
| Offspring only | Downregulated | 1669 | 8.38 (4.17) |
| Both | Upregulated | 19 | 0.01 (0.11) |
| Both | Downregulated | 112 | 0.01 (0.11) |
| Parental: Upregulated<br>Offspring: Downregulated | Opposite | 7 | 0.01 (0.11) |
| Parental: Downregulated<br>Offspring: Upregulated | Opposite | 30 | 0.01 (0.10) |

**Table S5:** Effects on offspring (F1) development time (days from oviposition to hatching). Maternal age (agetreat) was modelled as a fixed predictor and isoline identity was modelled as a random effect with gaussian error using glmmTMB. Dispersion was modelled as ~0 + agetreat. The data consisted of a total of 173 replicates from 10 isolines.

| | Estimate (conditional) | Std. Error | z-value | p-value |
|---|---|---|---|---|
| Intercept | 7.120 | 0.249 | 28.607 | < 0.001 |
| Maternal age treatment | 0.068 | 0.313 | 0.219 | 0.827 |

**Table S6:** Effects on offspring (F1) age at first reproduction (days from hatching to first eggs laid). Maternal age (agetreat) was modelled as a fixed predictor and isoline identity was modelled as a random effect with negative binomial error with quadratic parameterization using glmmTMB. Dispersion was modelled as ~0 + agetreat. The data consisted of a total of 180 replicates from 10 isolines.

| | Estimate (conditional) | Std. Error | z-value | p-value |
|---|---|---|---|---|
| Intercept | 2.857 | 0.0346 | 82.48 | < 0.001 |
| Maternal age treatment | -0.195 | 0.0407 | -4.79 | < 0.001 |

**Table S7:** Effects on offspring (F1) lifetime fecundity (up to age 167 days). Maternal age (agetreat) was modelled as a fixed predictor and isoline identity was modelled as a random effect with negative binomial error with quadratic parameterization using glmmTMB. Dispersion was modelled as ~0 + agetreat. The data consisted of a total of 180 replicates from 10 isolines.

| | Estimate (conditional) | Std. Error | z-value | p-value |
|---|---|---|---|---|
| Intercept | 7.118 | 0.045 | 157.60 | < 0.001 |
| Maternal age treatment | 0.363 | 0.095 | 3.82 | < 0.001 |

**Table S8:** Effects on offspring (F1) lifetime fecundity (up to age 167 days), accounting for reproductive lifespan and mortality. Maternal age (agetreat), offspring reproductive lifespan (replifespan = days between the replicate's first and last reproduction) and the mean number of individuals alive in the replicate over its reproductive lifespan (Meanalive) were modelled as fixed predictors, and isoline identity was modelled as a random effect with negative binomial error with quadratic parameterization using glmmTMB. Dispersion was modelled as ~0 + agetreat + replifespan + Meanalive. The data consisted of a total of 181 replicates from 10 isolines.

| | Estimate (conditional) | Std. Error | z-value | p-value |
|---|---|---|---|---|
| Intercept | 6.665 | 0.499 | 13.366 | < 0.001 |
| Maternal age treatment | 0.588 | 0.061 | 9.692 | < 0.001 |
| Reproductive lifespan | 0.005 | 0.002 | 2.616 | 0.0089 |
| Number alive | -0.045 | 0.091 | -0.495 | 0.6203 |

**Table S9:** Effects on offspring (F1) age-specific fecundity (up to age 167 days). Offspring age (Age), maternal age (agetreat), and the mean number of individuals alive in the replicate (Alive) were modelled as fixed predictors, and isoline identity and replicate identity were modelled as a random effects with negative binomial error with linear parameterization using glmmTMB. Dispersion was modelled as ~0 + Age + agetreat + Alive, and zero inflation was modeled as ~0 + Age * agetreat + Alive + (1|iso). The data consisted of a total of 932 observations (7-day fecundity totals per replicate) from 182 replicates and 10 isolines.

| | Estimate (conditional) | Std. Error | z-value | p-value |
|---|---|---|---|---|
| Intercept | 5.270 | 0.235 | 23.379 | < 0.001 |
| Age | -0.006 | 0.001 | -7.054 | < 0.001 |
| Maternal age treatment | 0.910 | 0.143 | 6.384 | < 0.001 |
| Number alive | 0.146 | 0.042 | 3.442 | < 0.001 |
| Age x Maternal age treatment | -0.006 | 0.002 | -3.784 | < 0.001 |

**Table S10:** Likelihood ratio test for the offspring age x maternal age interaction. A full model (see Table S9) was compared with an identical model lacking the interaction term (main-effects model).

| Model | Model DF | AIC | BIC | Log-likelihood | deviance | Chi-sq | Chi-sq DF | *p*-value |
|---|---|---|---|---|---|---|---|---|
| Full | 18 | 11424 | 11511 | -5694.1 | 11388 | | | |
| Main-effects | 17 | 11434 | 11516 | -5700.0 | 11400 | 11.791 | 1 | < 0.001 |

**Table S11:** Cox proportional hazards model comparing mortality rate of offspring of older mothers (OF) versus offspring of young mothers (OFY). Time to replicate extinction (defined as 2 or fewer individuals out of 5 left alive in the replicate) was fitted as a fixed predictor, and isoline identity was modelled as a random effect. Replicates with > 2 individuals alive at age 167 days were modelled as censored data. The test is based on 182 replicates from 10 isolines. The effect of maternal age at reproduction was tested using a Wald test ($W = 54.08$, df= 1, $p < 0.001$).

| | Coef | exp(coef) | Std. Error | Robust Std. Error | *z*-value | *p*-value |
|---|---|---|---|---|---|---|
| Maternal age treatment | 1.9280 | 6.8755 | 0.2950 | 0.2622 | 7.354 | < 0.001 |

**Table S12:** Effects on offspring (F1) body size (total body length; Fig. S2). Treatment (PAY, PAO, OFY, OFO) was modelled as a fixed predictor and isoline identity and replicate identity were modelled as random effects with gaussian error using glmmTMB. The data consisted of a total of 273 individuals from 30 replicates and 10 isolines.

| | Chi-sq | DF | p-value |
|---|---|---|---|
| Intercept | 1248.4 | 1 | < 0.001 |
| Treatment (PAY, PAO, OFY, OFO) | 326.8 | 3 | < 0.001 |

**Table S13:** Two-way contrasts between treatment groups based on Tukey tests for the offspring body size model (Table S12).

| Contrast | Estimate | Std. Error | *t*-value | *p*-value |
|---|---|---|---|---|
| OFO - OFY | -0.146 | 0.0334 | -4.383 | 0.0001 |
| OFO - PAO | -0.651 | 0.0378 | -17.245 | <0.0001 |
| OFO - PAY | -0.269 | 0.0362 | -7.414 | <0.0001 |
| OFY - PAO | -0.505 | 0.0351 | -14.396 | <0.0001 |
| OFY - PAY | -0.122 | 0.0340 | -3.596 | 0.0022 |
| PAO – PAY | 0.383 | 0.0374 | 10.249 | <0.0001 |

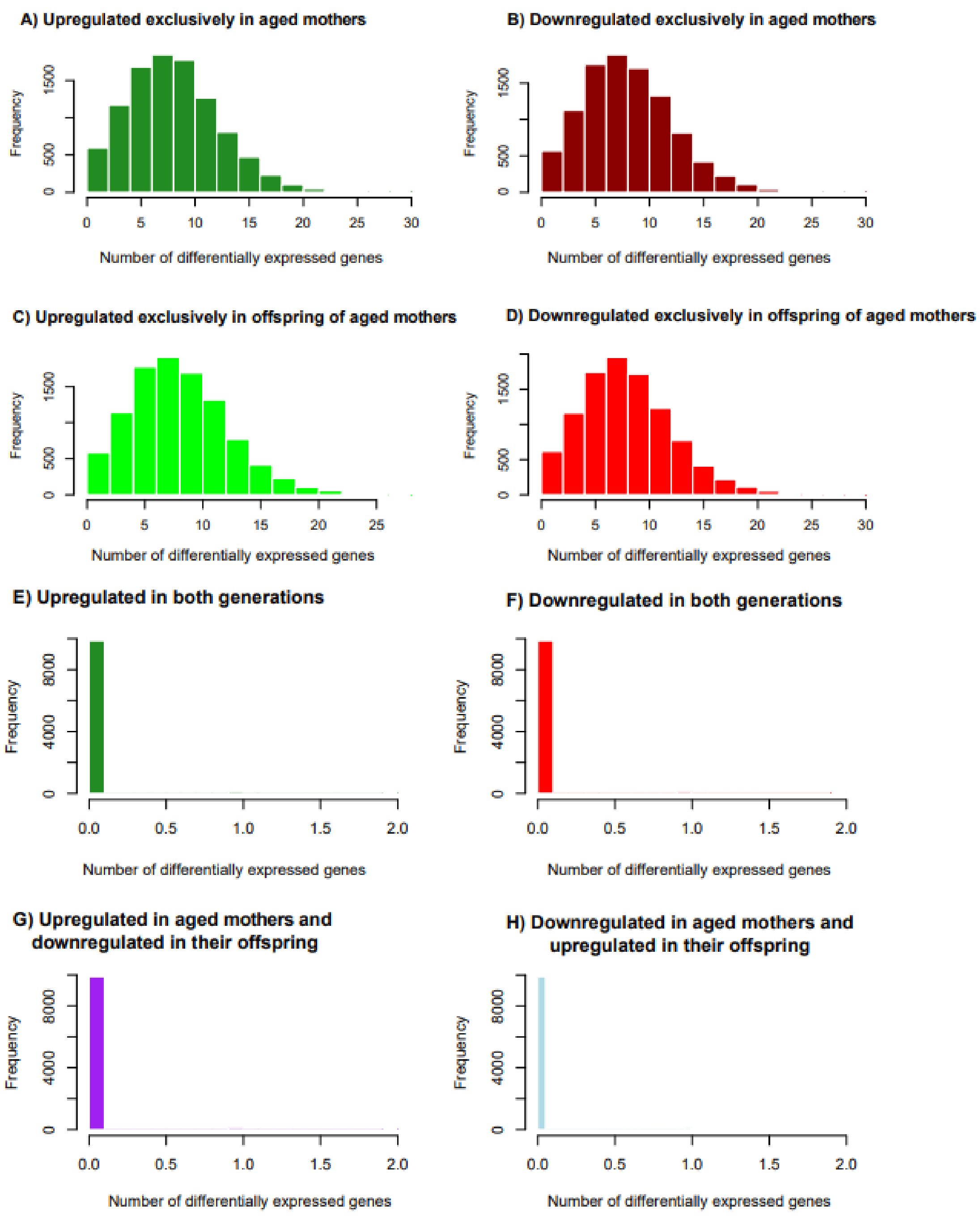


**Figure S2:** Randomization test results showing the expected number of differentially expressed genes under the null hypothesis. The histograms represent the distribution of the number of genes identified by *DESeq2* as differentially expressed (DEGs) across 10,000 randomization iterations for each category: (A) upregulated and (B) downregulated exclusively in aged mothers; (C) upregulated and (D) downregulated exclusively in offspring of aged mothers; (E) upregulated or downregulated in both generations; and (F) showing opposing regulation between generations. Observed values fall far outside the null distributions (see Table 3 in main manuscript), indicating non-random patterns of differential expression.

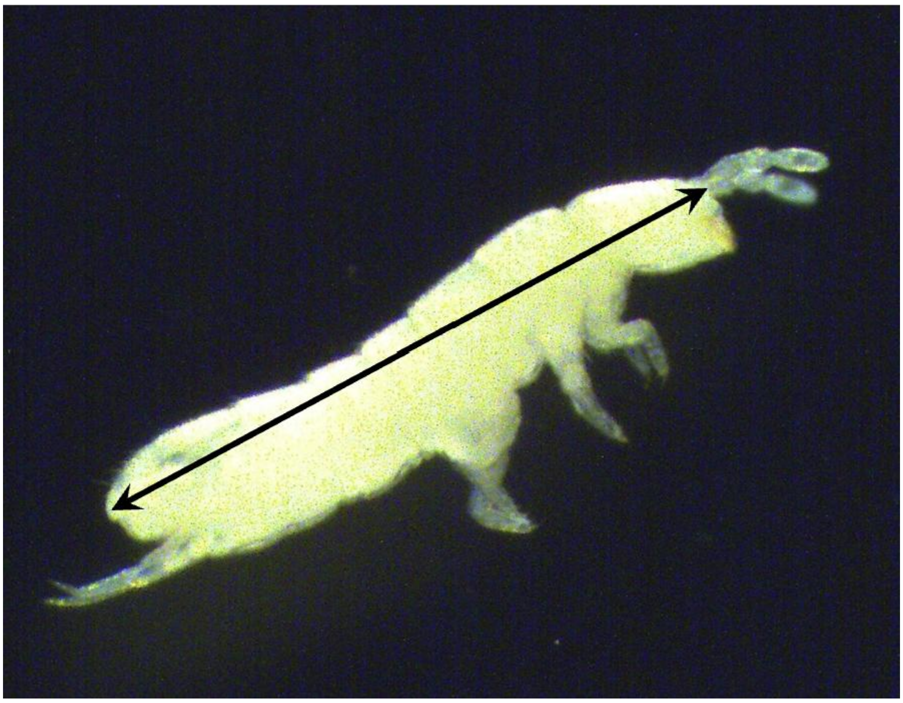

**Figure S3**. Maternal and hatchling body size was quantified from images taken with a Leica MC170 HD camera mounted on a Leica MZ 16A stereoscope. Total body length was measured from the posterior margin of the abdomen to the anterior margin of the head, not including the antennae using ImageJ software.

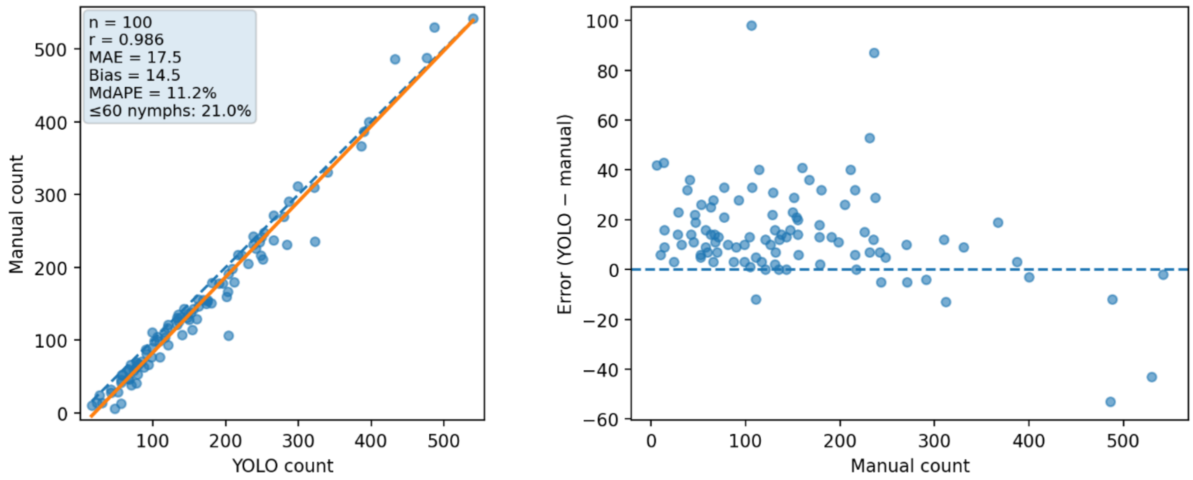


**Figure S4.** Validation of automated hatchling nymph counting. Automated counts generated by the trained YOLOv8 model were compared with manual counts across an independent validation set of 100 container images not used during model training. Left panel: The relationship between the automated (YOLO) and manual counts. Points represent container-level counts obtained by recombining detections from tiled image inference. Approximately 21% of validation images contained ≤60 individuals. The orange line indicates the fitted linear regression, while the dashed blue line shows a 1:1 relationship between automated and manual counts. Automated counts showed strong agreement with manual counts (Pearson r = 0.986; mean absolute error = 17.5 individuals). Right panel: Residual error (YOLO − manual) across the observed range of manual counts. Automated counts tended to over-estimate actual counts in images with low numbers of hatchlings. Correcting for this bias did not alter any results, so analyses based on uncorrected counts are reported. Code, trained model weights, and image-processing scripts are available on the Github repository (https://github.com/wyldescience/FolSum).

References


1. DeStefano, M. A. & Jacinto, E. Regulation of insulin receptor substrate-1 by mTORC2 (mammalian target of rapamycin complex 2). *Biochem. Soc. Trans.* **41**, 896–901 (2013).
2. Ghosh, H. S., McBurney, M. & Robbins, P. D. SIRT1 Negatively Regulates the Mammalian Target of Rapamycin. *PLoS ONE* **5**, e9199 (2010).
3. Smiles, W. J. *et al.* New developments in AMPK and mTORC1 cross-talk. *Essays Biochem.* **68**, 321–336 (2024).
4. Wek, R. C. Role of eIF2α Kinases in Translational Control and Adaptation to Cellular Stress. *Cold Spring Harb. Perspect. Biol.* **10**, a032870 (2018).
5. Proud, C. G. mTORC1 signalling and mRNA translation. *Biochem. Soc. Trans.* **37**, 227–231 (2009).
6. Wang, L., Lawrence, J. C., Sturgill, T. W. & Harris, T. E. Mammalian target of rapamycin complex 1 (mTORC1) activity is associated with phosphorylation of raptor by mTOR. *J. Biol. Chem.* **284**, 14693–14697 (2009).
7. Dunkerly-Eyring, B. L. *et al.* Single serine on TSC2 exerts biased control over mTORC1 activation mediated by ERK1/2 but not Akt. *Life Sci. Alliance* **5**, e202101169 (2022).
8. Kim, L. C., Cook, R. S. & Chen, J. mTORC1 and mTORC2 in cancer and the tumor microenvironment. *Oncogene* **36**, 2191–2201 (2017).
9. Simonson, B. *et al.* DDiT4L promotes autophagy and inhibits pathological cardiac hypertrophy in response to stress. *Sci. Signal.* **10**, eaaf5967 (2017).
10. Wen, J.-K. *et al.* Atg9 antagonizes TOR signaling to regulate intestinal cell growth and epithelial homeostasis in Drosophila. *eLife* **6**, e29338 (2017).
